# Benchmarking 80 binary phenotypes from the openSNP dataset using deep learning algorithms and polygenic risk score tools

**DOI:** 10.64898/2026.03.06.710126

**Authors:** Muhammad Muneeb, David B. Ascher, YooChan Myung, Samuel F. Feng, Andreas Henschel

**Affiliations:** School of Chemistry and Molecular Biology, The University of Queensland, Queen Street, 4067, Queensland, Australia; Computational Biology and Clinical Informatics, Baker Heart and Diabetes Institute, Commercial Road, 3004, Victoria, Australia; Department of Science and Engineering, Sorbonne University Abu Dhabi, Hazza Bin Zayed St, 38044, Abu Dhabi, United Arab Emirates; Department of Electrical Engineering and Computer Science, Khalifa University of Science and Technology, Saada Street, 127788, Abu Dhabi, United Arab Emirates

**Keywords:** genotype-phenotype prediction, machine learning, deep learning, openSNP, polygenic risk scores

## Abstract

Genotype-phenotype prediction plays a crucial role in identifying disease-causing single nucleotide polymorphisms and precision medicine. In this manuscript, we benchmark the performance of various machine/deep learning algorithms and polygenic risk score tools on 80 binary phenotypes extracted from the openSNP dataset. After cleaning and extraction, the genotype data for each phenotype is passed to PLINK for quality control, after which it is transformed separately for each of the considered tools/algorithms. To compute polygenic risk scores, we used the quality control measures for the test data and the genome-wide association studies summary statistic file, along with various combinations of clumping and pruning. For the machine learning algorithms, we used p-value thresholding on the training data to select the single nucleotide polymorphisms, and the resulting data was passed to the algorithm. Our results report the average 5-fold Area Under the Curve (AUC) for 29 machine learning algorithms, 80 deep learning algorithms, and 3 polygenic risk scores tools with 675 different clumping and pruning parameters. Machine learning outperformed for 44 phenotypes, while polygenic risk score tools excelled for 36 phenotypes. The results give us valuable insights into which techniques tend to perform better for certain phenotypes compared to more traditional polygenic risk scores tools.

## Introduction

The interactions of Deoxyribonucleic Acid (DNA) and environmental factors determine humans’ physical characteristics [1, 2]. The ability to determine phenotypes, especially disease risk, using genomic data, is increasingly important in the era of precise medicine [3, 4]. A genetic variation or difference occurring in DNA at a single position, where one nucleotide (A, T, C, or G) replaces another, is called single-nucleotide polymorphism (SNP). The identification of SNPs in the DNA responsible for the variation in phenotypes can help drive new diagnostic and therapeutic options [5, 6].

There are four primary methods used for genotype-phenotype prediction. The first is Genome-wide association studies (GWAS) [7, 8], in which the large-scale genotype data is analyzed to find associations between genetic variants and complex traits. GWAS can miss rare mutations or gene-gene relationships. It is limited to population-level associations and requires large sample sizes [9].

The second is Polygenic risk scores (PRS) [10, 11], a numerical score used in genetics to estimate an individual’s genetic predisposition for a particular trait, disease, or outcome. PRS is calculated by considering a person’s genetic variants, mainly single nucleotide polymorphisms (SNPs), and their associated weights or effect sizes (calculated from the GWAS). Such models have been used for the prediction of diseases and phenotypes [12], which include Type-2 diabetes [13], Hyperactivity Disorder [14], and Cardiovascular Disease [15]. The third is Computational modeling [16, 17, 18], in which genetic variation’s effect on the biological process is studied. Computational models can integrate various data types (genomics, transcriptomics, proteomics, and environmental factors). By integrating these data sources, models can better capture complex interactions and provide a deeper comprehension of the genes, regulatory pathways, and gene-gene interactions that contribute to variation in phenotype.

The fourth is machine/deep learning (ML/DL) [19, 20, 21, 22, 23, 24], in which genotype data is passed to a model for case/control classification. These methods can handle large datasets, capture non-linear interactions among SNPs in high-dimensional feature spaces, and integrate several genomic data types (genotype and gene data) [25] to enhance prediction. The ML/DL pipelines [26, 27, 28, 29] for genotype-phenotype prediction have been used for humans (facial features [30, 31], diseases [32], and behavioral phenotypes [33]), plants [34, 35, 36, 37, 38], and animals. Researchers used multiple (Random forest, SVM, gradient boosting machines) ML algorithms [39], L1L2 regularization [40], deep convolution neural networks [41], multilayer perceptrons [42], and combination of random forest and SVM [43] for genotype-phenotype prediction. We included most of these algorithms in our study. Apart from the methods mentioned above, genomic selection [44], transcriptomics and proteomics [45], and phenome-wide association studies [46] can be used for genotype-phenotype prediction.

Among publicly available datasets containing volunteer data is openSNP, which we used in this manuscript for analysis. This dataset contains multiple phenotypes and genotype data that are sufficient for PRS and ML/DL analysis. The limited number of samples for each phenotype made it possible to test multiple ML/DL hyperparameters and PRS clumping and pruning parameters for case-control classification [47]. Such efforts to explore the genotype-phenotype prediction using openSNP data have been made by multiple studies.

Naret et al. [48] calculated PRS for height using ML techniques on genotype data from openSNP and achieved an explained variance of 0.53, indicating that the PRS model explained 53 percent variation in height using genotype data. Saha et al. [49] used graph theory to find multi-loci associated with Astigmatism. They replaced the p-value thresholding with graph theory to select SNPs. Rajesh et al. [50] used statistical approaches and ML to predict diabetic macular oedema from openSNP data and achieved an F1 score = 0.84, indicating that the model performed well in distinguishing cases and controls. Lu et al. [51] used a Bayesian Network for SNPs and trait inferences for multiple traits using data from openSNP. Javier et al. [52] proposed a tool for cases/controls classification in R language, which uses eight ML models for benchmarking. Muneeb et al. [53] used ML/DL models for eye color and type-2 diabetes prediction using openSNP data. In all the studies, researchers considered at most ten phenotypes from openSNP for SNPs and trait correlation analysis. The phenotype transformation process, the number of samples they considered, and the ML/DL models they used differ significantly from our study.

In this work, we benchmarked the performance of 29 ML algorithms, 80 DL algorithms variants, and 3 PRS tools with 675 clumping and pruning parameters on 80 phenotypes from the openSNP dataset [54]. The comparison of ML and PRS tools is noteworthy, as for some phenotypes (Parkinson’s Disease [55] and Breast cancer [56]), ML/DL algorithms produced better results. On the other hand, PRS outperforms ML for some phenotypes (coronary artery disease [57]). Genotype data is typically extensive, and deep learning algorithms can process large datasets (through batch processing) compared to classical machine learning algorithms, which require computational resources and much time to execute. The deep learning approaches are often combined with PRS to increase the classification performance [58, 59, 60, 61]. In this study, ML/DL algorithms performed better for 44 phenotypes and PRS tools for 36 phenotypes.

Genotype-phenotype prediction offers various benefits beyond identifying disease-associated SNPs, including disease risk prediction, early disease prediction, and case-control classification for screening. Our study primarily focuses on case-control classification rather than the identification of specific SNPs [62]. The latter requires extensive data and various statistical measures to establish associations. Below, we have enumerated the research contributions that have given shape to novel findings and research questions, serving as the driving force behind this research work.

Ma et al. [63] concluded that different PRS methods exhibit distinct performance characteristics across traits. Moreover, the same method may exhibit varying performance when applied to the same trait in different studies, owing to differences in cross-validation designs. The performance of the best-performing model depends on the p-value threshold, clumping, and pruning parameters. These parameters can vary across populations, datasets, and phenotypes being considered. Similarly, hyperparameters in ML and DL algorithms impact classification performance. We also compared and unified the steps required by both pipelines, shedding light on the processes where methodologies differ between using PRS [10] and ML/DL [26] for case-control classification.

Our research also involved the study of phenotypes, physical characteristics, appearance, and preferences, for which genetic causes may not be apparent. The model’s performance was notably low for such phenotypes, including preferences such as enjoying driving a motorbike, sports interests, and fishing. This suggests that these preferences are more likely results of environmental factors rather than genetic influence, which is consistent with observations in a study by English et al. [64].

Transfer learning is a widely used technique in genetics [65, 66]. Models trained on extensive data from a well-studied, large population are utilized for case-control classification in smaller, understudied populations with limited data. Our research explores how modifications in hyperparameters can lead to improved baseline performance in smaller populations [67].

## Methodology

Data, data transformation and modeling, quality control steps on genotype data, ML/DL techniques, and PRS tools used for case-control classification are explained in the methodology. Figure 1 shows a graphical representation of the methodology presented in this study.

**Fig. 1.**
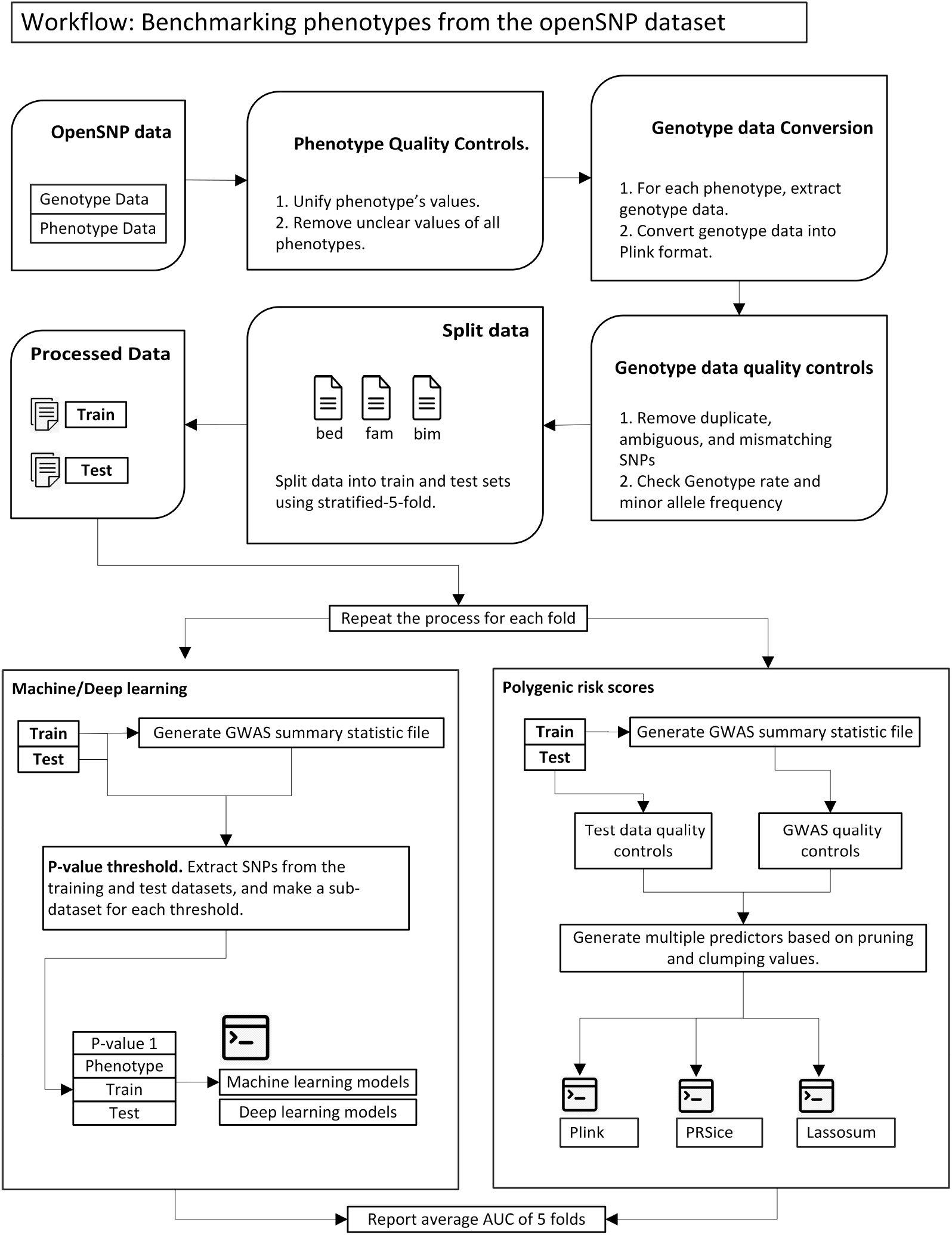
A workflow of genotype-phenotype prediction using ML/DL and PRS. A case/control classification flowchart using ML/DL and PRS tools. First, clean phenotype data and extract binary phenotypes from the openSNP dataset. Second, merge the genotype data for each phenotype, convert the dataset to Plink, and perform quality controls on genotype data. Split data in train/test and the processing is done on each fold. Third, generate multiple sub-datasets using p-value thresholds on the GWAS file, and pass the sub-datasets to ML/DL models. Fourth, generate the GWAS from the training data and perform quality controls on both test and GWAS files. After that, the processed GWAS and test are passed to PRS tools for processing.

We considered binary phenotypes from the openSNP dataset. We cleaned the phenotype data because it contained inconsistent values for each phenotype, and we unified varying values into two phenotype values to generate cases and controls for each phenotype. This process is referred to as phenotype cleaning or transformation [68], and it is a manual and time-consuming process. We converted all files in AncestryDNA and 23andme format to Plink file format (bed, bim, fam) [69]. The detailed data pre-processing steps are explained in the Supplementary material, Section 1.

### Modeling

After data pre-processing, we had 101 phenotypes in Plink file format, and the next step was to improve the dataset quality. We considered a minor allele frequency = 0.01, Hardy-Weinberg equilibrium = 1e-6, genotype rate = 0.01, and missing individuals rate = 0.7. At this stage, duplicate files for each person and duplicate SNPs were removed. After this process, we had 80 phenotypes, and a list is available on GitHub (Analysis3.pdf).

We used Plink to split (stratified 5-fold) the data into training (80%) and test sets (20%), which can be done by dividing the phenotype file into two sets and extracting the subset of people belonging to each set. After this step, we performed the following steps on each fold. In the following text, the ML/DL analysis is described in section 2.1.1 and for PRS in section 2.1.2.

#### Machine/Deep learning - Modeling

The genetic data usually consists of millions of SNPs (features), and using p-value thresholding to reduce the number of SNPs before ML/DL models is a regular procedure for genotype-phenotype prediction [70, 71, 72]. A GWAS summary statistic file is required for p-value thresholding. We subjected the training data to Fisher’s exact test for allelic association (using `plink --assoc`) to generate the GWAS summary statistics containing p-values for each SNP. For ML and DL, we extracted 50, 100, 200, 500, 1000, 5000, and 10000 SNPs from the training and test data, saving the encoded data in raw spreadsheet format to train the machine and deep learning algorithms. The GWAS file contains p-values for SNPs, and we linearly increased the p-value threshold (starting from 0) to extract a specific number of SNPs. A specific p-value threshold required to extract a particular number of SNPs can vary among different phenotypes

Silva et al. [73] and Medvedev et al. [74] used machine learning models for SNP selection and genotype-phenotype prediction. One potential reason for this is the ability of machine and deep learning models to consider highly explanatory variables during the training phase.

The next step was training the ML/DL model and testing the performance. We used the default parameters for the ML models and mutated a few hyperparameters for the DL models. We changed the model’s hyperparameter to get a new variant of a specific model. Finally, we averaged the Area under the receiver operating curve (AUC) of all folds and reported the model’s performance and corresponding hyperparameter [75], which yielded the best AUC.

#### PRS - Modeling

PRS tools require two files: Base file and Target files. The base file is a GWAS summary statistic containing p-values and odds ratios for each SNP, generated using the training data. GWAS summary statistic file should be in a particular format and require extra steps so PRS tools like Plink, PRSice, and Lassosum can process it. We generated the GWAS data from the training dataset rather than relying on existing studies. It’s important to note that the GWAS file is available for only 30 out of the 80 phenotypes. Furthermore, the existing GWAS data primarily represents individuals with European ancestry. In contrast, openSNP respondents predominantly come from the USA (60.33%), followed by Canada (5.17%) and the UK (4.61%). The genotype rate in openSNP is lower compared to the genotype rate in the GWAS data from existing research projects. This discrepancy may result in PRS scores based on a limited number of SNPs. To address this, we generated the GWAS data from our target dataset. It’s worth mentioning that the number of samples available for most phenotypes is relatively low but still meets the minimum criteria, as specified by Crossa et al. [76] and Qu et al. [77].

The list of columns in the GWAS summary required by each PRS tool is mentioned in their corresponding documentation and this tutorial https://choishingwan.github.io/PRS-Tutorial/base/. Chromosomes (CHR), Base pair (BP), SNP ID (SNP), p-value (P), and effect size/odds ratio (OR) are generated using Plink (Fisher’s exact test). The standard error of association (SE) was calculated using P and OR. Reference allele (A1), Alternative allele (A2), Minor allele frequency (MAF), and the total number of people (N) are generated using SNPTEST. Finally, we merged the information from both GWAS files to generate the final GWAS file, as required by PRS tools. The PRS tools we considered require CHR, BP, SNP, P, OR, A1, A2, N, and MAF to generate the PRS score. We generated the exact GWAS file with the same headers, simplifying the following calculation.

PRS calculation requires quality control steps on the GWAS file and test data before the PRS tool can process the information. The quality control steps for GWAS data involved removing duplicate, ambiguous, and mismatched SNPs. We considered SNPs having imputation information score *>* 0.8 and minor allele frequency *>* 0.01. The quality control steps for the target data involved: removing duplicate, ambiguous, mismatching SNPs. We considered SNPs with a Genotype rate *>* 0.01, minor allele frequency *>* 0.01, and hardy-Weinberg equilibrium = 1e-6. Individuals having a missing rate per SNP below 0.7 were removed from the analysis. The rest of the analysis is performed on the common SNPs between the GWAS and target files [78, 79]. An explicit p-value threshold is not required for PRS tools, as they automatically increase the p-value threshold from minimum to maximum with a particular interval to find the optimal p-value threshold. Pruning involves searching through genomes for linked SNP pairs and keeping only the SNP with the highest minor allele frequency. Clumping is used to identify genetic variants that are highly correlated with each other and potentially associated with a particular trait or disease [80, 67]. We considered multiple values of pruning and clumping on test data before it is passed to PRS tools, and those values are shown in Table 2.

The final step was to convert the PRS to AUC to compare the results with the ML models. The final layer of the deep learning model employs a sigmoid function to convert values to 0 or 1, typically using a threshold of 0.5. We then converted the final PRS scores to binary values (0 and 1) and compared them with the actual values for evaluation. To ensure consistency, we normalized the PRS scores for all individuals using Min-Max normalization, scaling them between 0 and 1. Individuals with PRS scores *<* 0.5 were classified as controls, while those with PRS scores *>* 0.5 were classified as cases. It is crucial to maintain a single threshold value across all folds when calculating PRS.

### Implementation

#### Machine learning - models

We used classical ML algorithms with default parameters from `scikit-learn` [81] library. The algorithms include tree-based classification algorithms [82] (AdaBoost, XGBoost [83], Random Forest, and Gradient boosting algorithms), SGD (Stochastic Gradient Descent), MLP (Multi-layer perceptron), SVC (Support Vector Classifier [84]), and other algorithms. The list of 29 ML algorithms and corresponding hyper-parameters are available on GitHub (MachineLearningAlgorithms.txt).

#### Deep learning - models

We used four basic DL models: Artificial neural network (ANN) [85], Gated recurrent unit (GRU) [86], Long Short-Term Memory (LSTM) [87], and (Bidirectional LSTM) BILSTM [88]. Recurrent neural networks were used for genotype-phenotype prediction Pouladi et al. [89] and Srinivasu et al. [90]. The extracted SNPs from the GWAS file can be treated as a sequence of SNPs with high potential for association. After experimenting with various architectures, including different numbers of layers and neurons, we decided to use a neural network with five layers, which was also recommended by Uzair et al. [91] and Pérez-Enciso et al. [92]. The numbers of neurons in each layer were 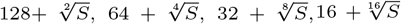, and 1, where S is the number of SNPs in the data. Genotype data comprises varying numbers of SNPs (features), necessitating the model to adapt to the input size. We explored three architectures: one with a fixed number of neurons, one with the input size halved, and one with the square root of the input size. The last architecture demonstrated superior performance. Consequently, we used the square root of the number of SNPs to determine the number of neurons at each layer. This approach allowed us to use the same model for sub-datasets of different dimensionalities. We created six new stacked model architectures using GRU, LSTM, and BILSTM layers. We generated 80 deep-learning models by changing the values of three hyperparameters and considering Adam as an optimizer, as shown in Table 1. This combination of hyperparameters resulted in 8 unique models of a specific architecture. The list of 80 deep learning algorithms is available on GitHub (DeepLearningAlgorithms.txt).

**Table 1.** Deep learning model’s hyperparameters and values. The first column shows the DL model’s hyperparameter, and the second column shows the values we considered for each hyperparameter. The combination of hyperparameter values resulted in 8 (2 ∗ 1 ∗ 2 ∗ 2) models for a specific architecture.

| Parameters | Values |
| --- | --- |
| Dropout | 0.2,0.5 |
| Optimizer | Adam |
| Batch size | 1,5 |
| Epochs number | 50,200 |

#### PRS tools

In the study by Choi et al., [10] the authors used Plink, PRSice2, Lassosum, and LDpred2 for PRS calculation. LDpred2, however, has some limitations. It utilizes SNPs from either the 1000 Genome project or HapMap3 SNPs and focuses on further analyzing common SNPs shared between the 1000 Genome project and the GWAS file. This approach is computationally demanding and restricts the ability to test multiple hyperparameters. Additionally, LDpred2 relies on heritability as a prerequisite for PRS calculation, which might not be suitable for phenotypes lacking genetic correlations. Consequently, we opted to use Plink [93], PRSice [94], and Lassosum [95] for our analysis.

Plink (Whole-genome association analysis toolset) is a widely used tool for working with genetic data, particularly in genome-wide association studies (GWAS). It provides a wide range of functions for quality control, data management, statistical analysis, and data visualization of genetic data. PRSice2 is a tool designed for the calculation and evaluation of PRS from genetic data. It constructs PRS by integrating information from multiple genetic variants and their effect sizes. Lassosum is a statistical method used to compute PRS by incorporating a LASSO (Least Absolute Shrinkage and Selection Operator) penalty. It aims to enhance PRS construction by selecting a subset of informative genetic variants while reducing the effect sizes of others to zero.

Once the quality control steps were performed on the GWAS summary statistic file and test set, the dataset was passed to PRS tools. Each tool uses a different formula for the calculation. After quality control steps, we performed pruning and clumping on the test data. Table 2 shows pruning and clumping variables and their values. The list of 675 PRS predictors is available on GitHub (Plink PRSice Lassosum Parameters.txt).

**Table 2.** PRS tools parameters and values. The first column shows the clumping and pruning parameters, and the second column shows the PRS values. Clumping p1 means a significant threshold for index SNPs, clumping r2 means linkage disequilibrium threshold for clumping, and clumping kb means physical distance threshold. The combination of hyperparameters values resulted in 675 (3 ∗ 3 ∗ 3 ∗ 1 ∗ 5 ∗ 5) PRS predictors for each tool.

| Parameters | Values |
| --- | --- |
| Pruning window size | 200,500,1000 |
| Pruning window shift size | 50,100,150 |
| Pruning LD threshold | 0.1,0.3,0.5 |
| Clumping p1 | 1 |
| Clumping r2 | 0.1,0.3,0.5,0.7,0.9 |
| Clumping kb | 200,400,600,800,1000 |

Plink calculates the PRS by summing the products of the effect sizes (usually beta coefficients) of each SNP and their corresponding genotypes (0, 1, or 2 copies of the risk allele), as specified in equation 1.

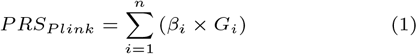

In equation 1, PRS means polygenic risk score, *β*_*i*_ represents the effect size or weight associated with the *i*th SNP (obtained from the GWAS file), and *G*_*i*_ means the genotype (or dosage) of the *i*th SNP with *G*_*i*_ ∈ {0, 1, 2}. *n* is the total number of SNPs. We used Plink (`plink --score`) to calculate PRS, and after that PRS score is compared with the actual phenotypes. Genotype data, PRS scores, and other covariates like age, gender, and the first six principal components can be used to retrieve explained variance by a linear model [10].

The Lassosum calculates the effect sizes of genetic variations using LASSO (Least Absolute Shrinkage and Selection Operator) regression and determines PRS by adding the predicted effect sizes and the matching genotypes for each SNP, as shown in equation 2. The idea of regularization is to reduce features contributing very little such that their weights are reduced to 0.

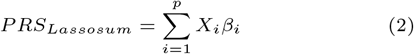

In equation 2, PRS represents Polygenic Risk Score, *β*_0_, *β*_1_, *β*_2_, …, *β*_*p*_ are the estimated coefficients, *X*_1_, *X*_2_, …, *X*_*p*_ represent SNPs included in the PRS calculation, and *p* is the number of SNPs.

The effect sizes of genetic variations are calculated using equation 3.

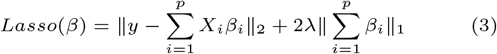

In equation 3, Lasso represents objective function, *β* are the coefficients to be determined by Lasso, *β*_0_, *β*_1_, *β*_2_, …, *β*_*p*_ are the coefficients to be estimated, *X*_1_, *X*_2_, …, *X*_*p*_ represent SNPs included in the PRS calculation, *λ* is the LASSO regularization parameter that shrinks coefficients towards zero and thus reduces overfitting, *y* is the known phenotype ground truth, and *p* is the number of SNPs.

PRSice offers four formulas to calculate the PRS. We used the default formula as specified in equation 4. In PRSice2, the PRS is calculated by summing the products of the estimated effect sizes and the corresponding genotypes, then dividing the sum by the number of SNPs.

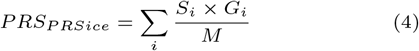

In equation 4, *i* represents a number of SNP, *S*_*i*_ and *G*_*i*_ represent the effect size and genotype value of *i*th SNPs, and *M* represents the number of SNPs.

## Results

The samples for most phenotypes were imbalanced, so we chose to consider AUC for reporting the classification performance, as recommended by multiple studies [96, 97, 74].

### Machine and deep learning results

First, we analyzed the results from ML/DL algorithms. Figure 2 shows the results from the ML/DL algorithms and groups them based on the SNPs that yielded the best results, as it helps to understand the complexity of the phenotypes. We considered seven p-value thresholds for each phenotype resulting in 7 sub-datasets containing 50, 100, 200, 500, 1000, 5000, and 10000. The optimal result for each phenotype should be from one of those p-value thresholds, and we grouped the phenotypes based on that threshold or the number of SNPs. The actual SNPs for each phenotype and p-value threshold are different.

**Fig. 2.**
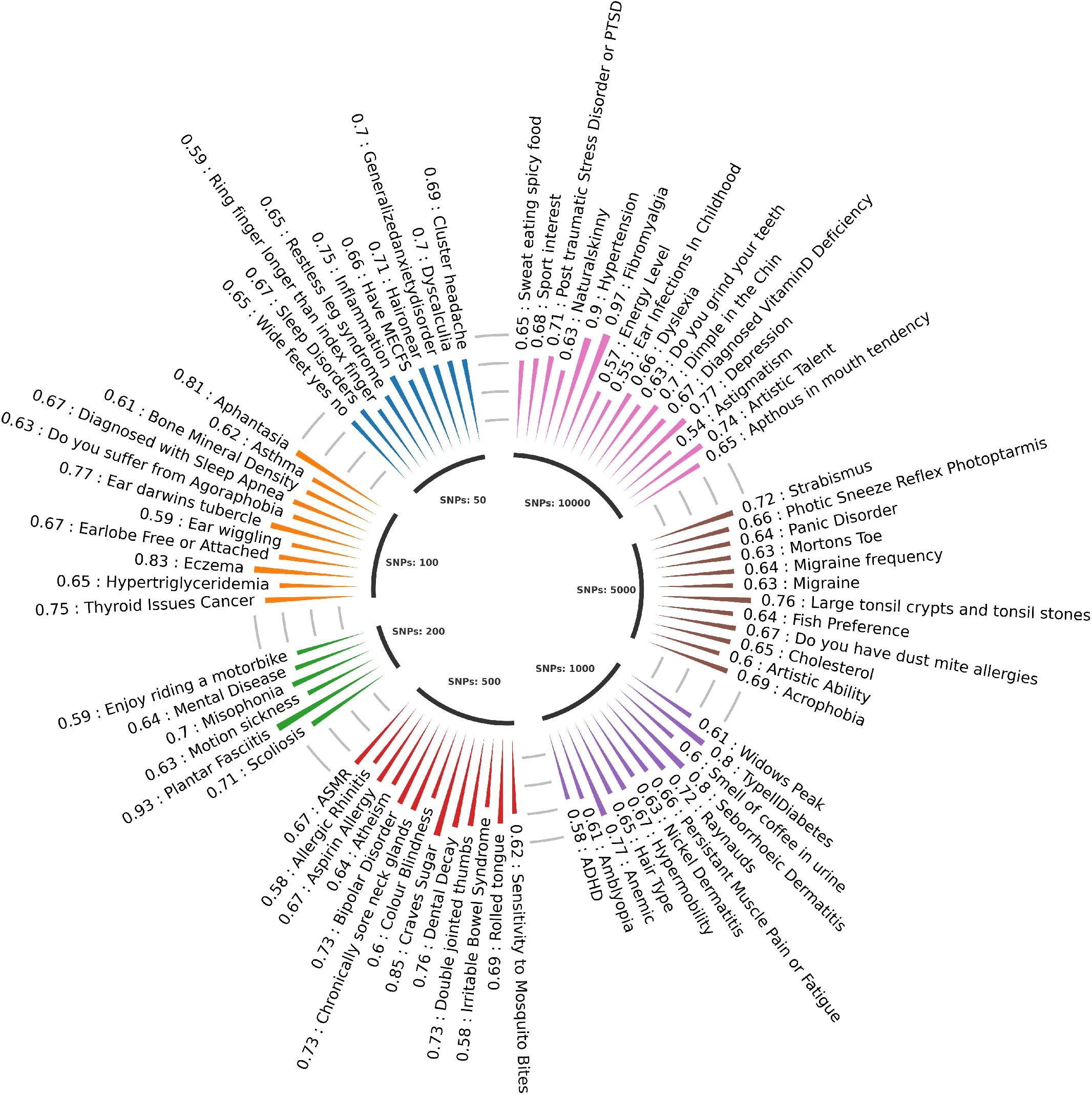
This diagram shows the AUC for each phenotype obtained from the ML/DL algorithms and group phenotypes on the number of SNPs that yield the best results.

For some phenotypes (Dyscalculia, Inflammation, and Restless leg syndrome), optimal results could be obtained by using a limited number of SNPs extracted from p-value thresholding. Some phenotypes (Type-2 Diabetes, Migraine, and Depression) require thousands of SNPs for optimal performance, indicating that a specific phenotype is complex 2. Among all ML algorithms SGDClassifier, XGBoost, PassiveAggressiveClassifier, DecisionTreeClassifier, MLPClassifier, and HistGradientBoostingClassifier turn out to be the best algorithms for genotype data resulting in the best performance for 12, 10, 9, 7, 6, and 6 phenotypes, respectively. Table 3 ranks the DL algorithms based on the number of phenotypes for which a specific algorithm produced the best results. Among DL algorithms, ANN is the best algorithm yielding the best results for 26 phenotypes. LSTM yielded the best result for only one phenotype.

**Table 3.**
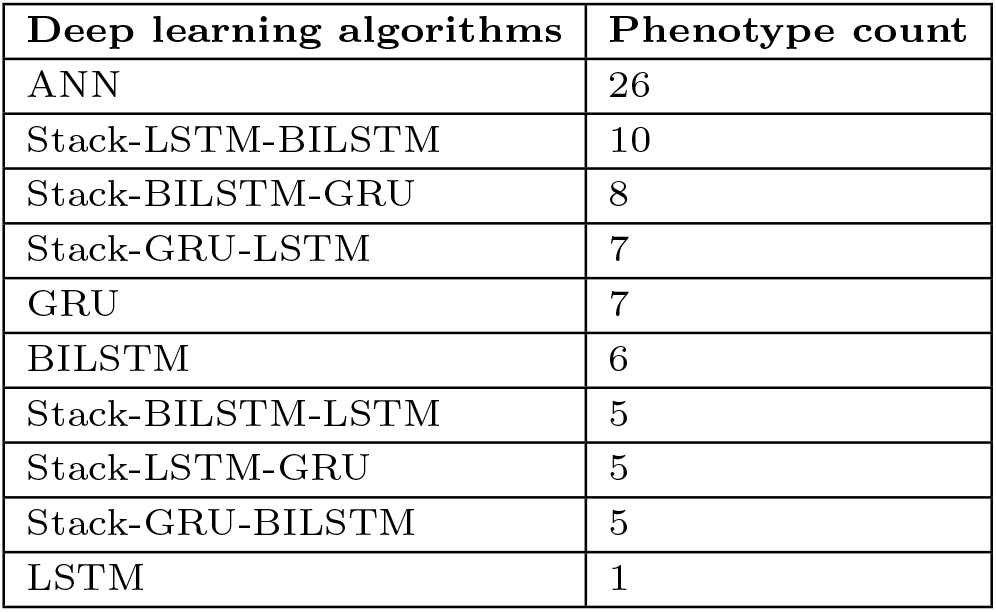
First column list the DL algorithm, and the second column shows the number of phenotypes for which the corresponding DL algorithm yielded the best results.

| Deep learning algorithms | Phenotype count |
| --- | --- |
| ANN | 26 |
| Stack-LSTM-BILSTM | 10 |
| Stack-BILSTM-GRU | 8 |
| Stack-GRU-LSTM | 7 |
| GRU | 7 |
| BILSTM | 6 |
| Stack-BILSTM-LSTM | 5 |
| Stack-LSTM-GRU | 5 |
| Stack-GRU-BILSTM | 5 |
| LSTM | 1 |

We changed values of three DL hyperparameters resulting in 8 variants, and Table 4 ranks the hyperparameters based on the number of phenotypes for which a specific hyperparameter combination resulted in the best performance. Dropout = 0.2, Optimizer= Adam, batch size = 1, and a number of epochs = 50 yielded the best performance for 23 phenotypes.

**Table 4.** First four columns list the DL algorithm parameter, and the second column (PC) shows the number of phenotypes for which the corresponding DL algorithm parameters yielded the best results. (D = Dropout, O = Optimizer, B = Batch size, and E = Number of Epochs).

| D | O | B | E | PC |
| --- | --- | --- | --- | --- |
| 0.2 | Adam | 1 | 50 | 23 |
| 0.2 | Adam | 5 | 50 | 18 |
| 0.5 | Adam | 5 | 50 | 16 |
| 0.5 | Adam | 1 | 50 | 15 |
| 0.2 | Adam | 5 | 200 | 4 |
| 0.5 | Adam | 1 | 200 | 2 |
| 0.2 | Adam | 1 | 200 | 1 |
| 0.5 | Adam | 5 | 200 | 1 |

### PRS tools results

We used 675 parameters for Plink as specified in Table 2. It can be seen in table 5 that mutations in pruning parameters did not improve the results, as no pruning parameter except pruning window size = 200, window shift size = 50, and pruning linkage disequilibrium threshold = 0.1 appeared in the best predictors. Mutations in clumping window size and linkage disequilibrium threshold improved the performance.

**Table 5.** The first six columns show the Plink parameters in this format (P1 = Pruning window size, P2 pruning window shift size, P3 pruning linkage disequilibrium threshold, C1 clumping window size, C2 = clumping p-value, C3 = clumping threshold). The second column (PC) shows the number of phenotypes for which specific predictors yielded the best results.

| P1 | P2 | P3 | C1 | C2 | C3 | PC |
| --- | --- | --- | --- | --- | --- | --- |
| 200 | 50 | 0.1 | 200 | 1 | 0.1 | 13 |
| 200 | 50 | 0.1 | 200 | 1 | 0.325 | 13 |
| 200 | 50 | 0.1 | 200 | 1 | 0.775 | 10 |
| 200 | 50 | 0.1 | 1000 | 1 | 0.1 | 8 |
| 200 | 50 | 0.1 | 400 | 1 | 0.1 | 5 |
| 200 | 50 | 0.1 | 400 | 1 | 0.775 | 4 |
| 200 | 50 | 0.1 | 200 | 1 | 0.55 | 4 |
| 200 | 50 | 0.1 | 1000 | 1 | 0.55 | 3 |

For PRSice, only one parameter combination (200-50-0.1-200-1-0.1) yielded the best results.See Table 2 for reference.

We used 675 parameters for Lassosum as specified in Table 2. It can be seen in Table 6 that mutations in clumping parameters did not improve the results, as no clumping parameter except clumping window size = 200, clumping p-value = 50, and clumping linkage disequilibrium threshold = 0.1 appeared in the best predictors. Mutations in pruning window size, pruning window shift size, and pruning linkage disequilibrium threshold improved the performance.

**Table 6.** The first six column show the Lassosum parameters in this format (P1 = Pruning window size, P2 = pruning window shift size, P3 = pruning linkage disequilibrium threshold, C1 = clumping window size, C2 = clumping p-value, C3 = clumping threshold). The second column (PC) shows the number of phenotypes for which specific predictors yielded the best results.

| P1 | P2 | P3 | C1 | C2 | C3 | PC |
| --- | --- | --- | --- | --- | --- | --- |
| 200 | 50 | 0.1 | 200 | 1 | 0.1 | 20 |
| 1000 | 50 | 0.1 | 200 | 1 | 0.1 | 8 |
| 500 | 50 | 0.3 | 200 | 1 | 0.1 | 5 |
| 200 | 100 | 0.3 | 200 | 1 | 0.1 | 4 |
| 1000 | 100 | 0.1 | 200 | 1 | 0.1 | 4 |
| 1000 | 100 | 0.3 | 200 | 1 | 0.1 | 4 |
| 1000 | 50 | 0.3 | 200 | 1 | 0.1 | 4 |
| 200 | 150 | 0.5 | 200 | 1 | 0.1 | 4 |
| 200 | 100 | 0.1 | 200 | 1 | 0.1 | 4 |

### Best-performing models

Figure 3 shows the heatmap produced by AUC from ML/DL algorithms, Plink, PRSice, and Lassosum. It can be seen that PRSice performs worst among all other approaches. ML/DL and Plink yielded better and comparable performance for each phenotype.

**Fig. 3.**
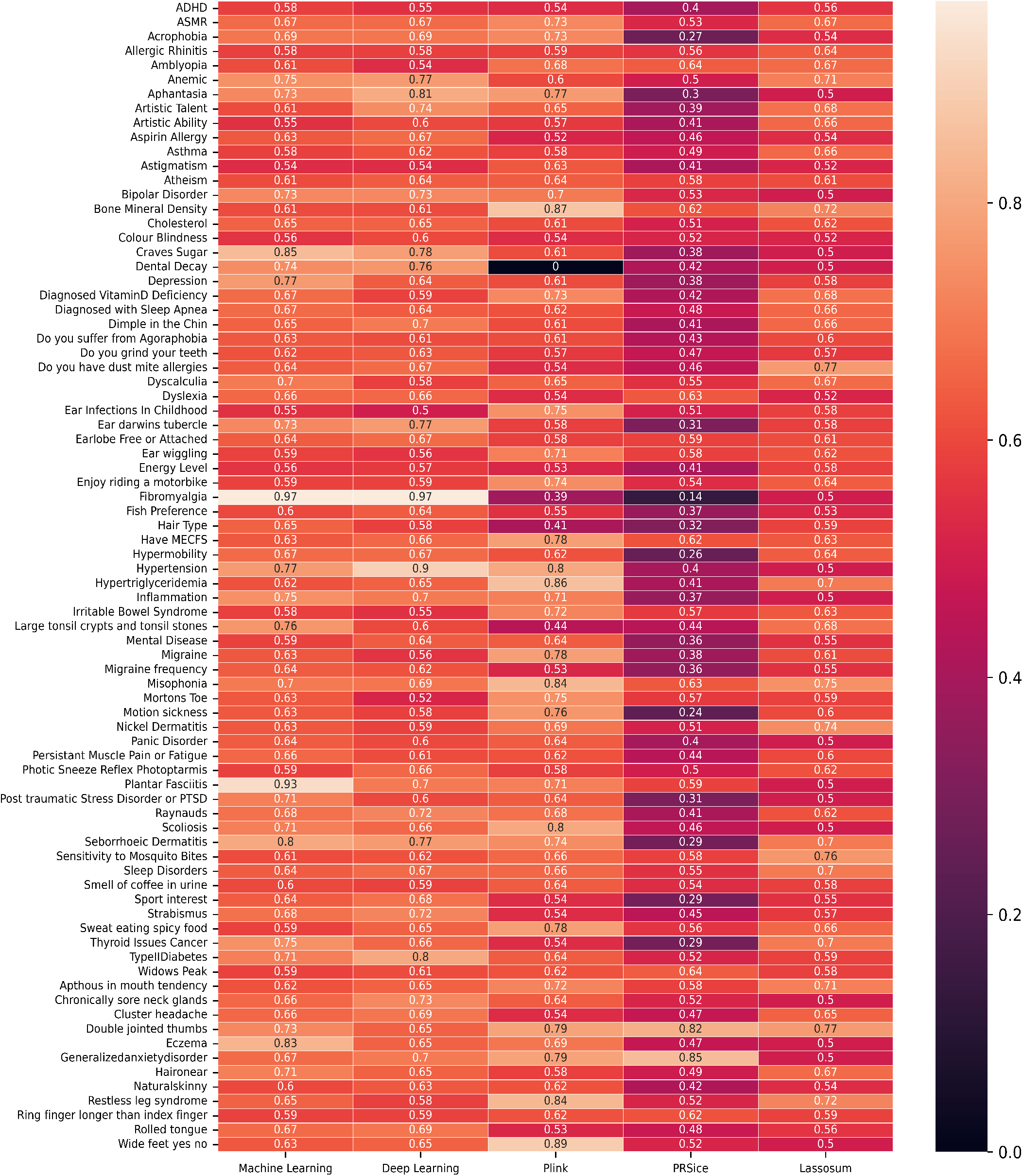
This heatmap shows the AUC as a percentage on the test data yielded by the ML/DL and PRS tools for each phenotype.

Figure 4 groups the phenotypes based on the tools which yielded the best AUC. It assists to understand whether ML/DL or PRS tools performed better for a specific phenotype. It can be seen that for 44 phenotypes, ML/DL achieved better performance. Among PRS tools, Plink produced better results. For TypeIIDiabetes, Seborrhoeic Dermatitis, Aphantasia, Eczema, Hypertension, Plantar Fasciitis, and Fibromyalgia ML/DL algorithms produced AUC ≥ 80 percent. For Scoliosis,

**Fig. 4.**
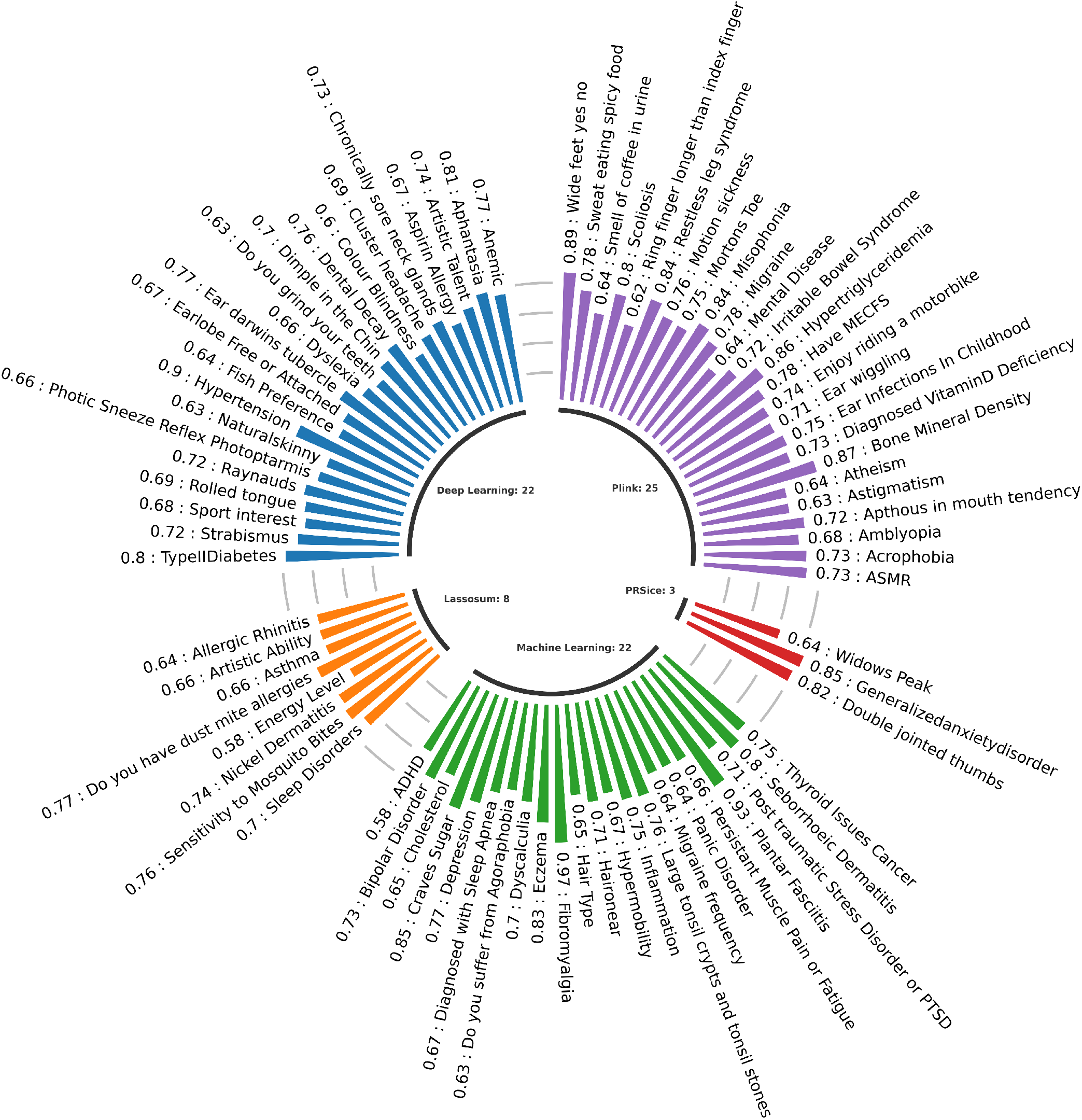
This diagram shows the best AUC for each phenotype and group phenotypes based on the tools that yield the best AUC for a specific phenotype.

Restless leg syndrome, Misophonia, Hypertriglyceridemia, and Bone Mineral Density PRS tools produced AUC ≥ 80 percent.

### Interpreting best-performing models

The genetic data quality and quantity, the underlying genetic architecture of phenotype, and the model used for classification impact the prediction’s performance. Moreover, hyperparameters and quality control steps like clumping, pruning, and p-value thresholding can significantly impact the performance, which is also evident in our results. We used 80 and 675 parameters for DL algorithms and PRS tools, so it is important to report the tool and hyperparameters or predictors that yielded the best AUC, as shown in Figure 5.

**Fig. 5.**
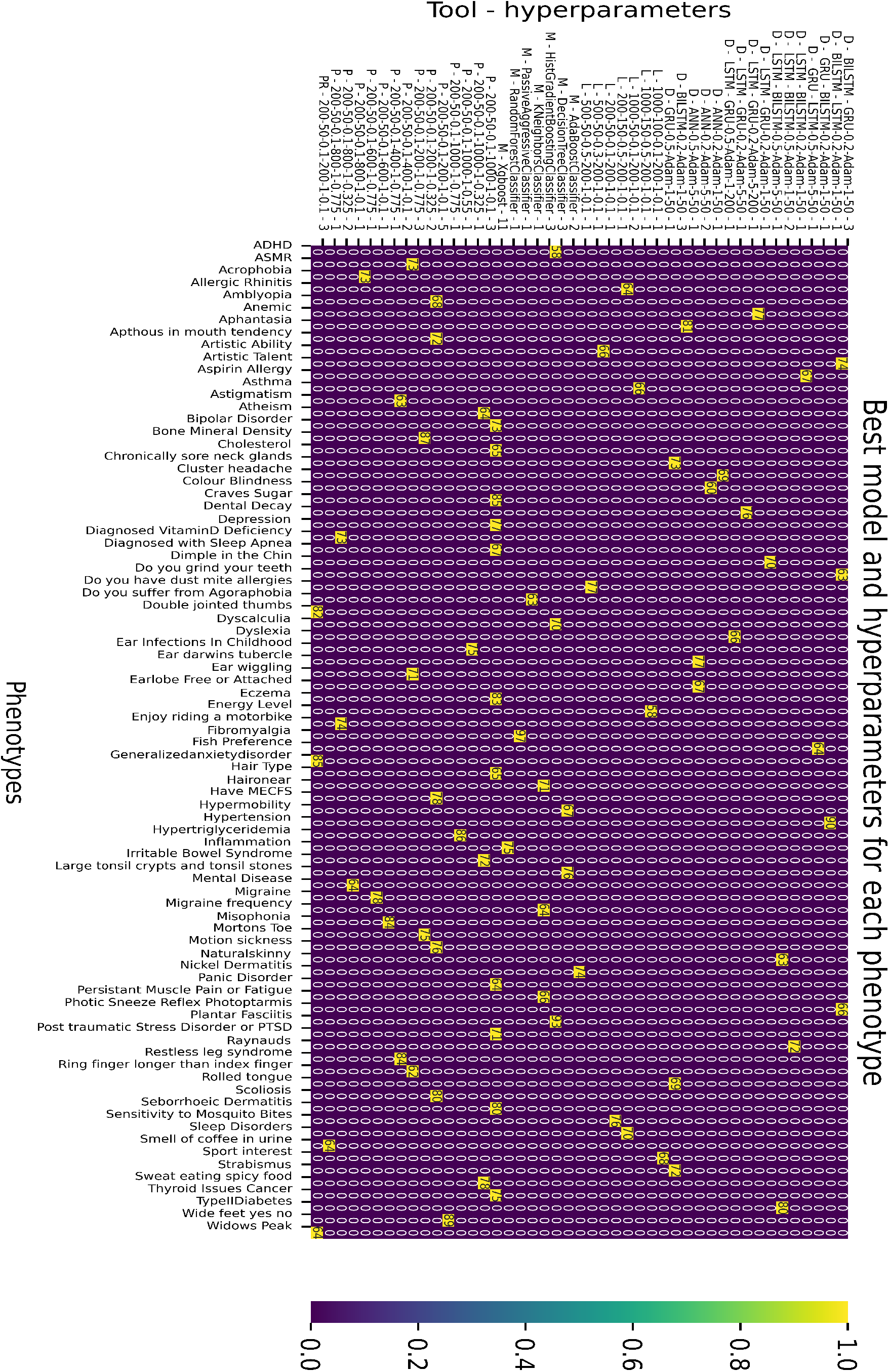
The x-axis shows a phenotype. The labels on the y-axis show the tool and hyperparameter that yielded the best AUC for a specific phenotype. D, L, M, P, and PR on y-axis mean Deep learning, Lassosum, Machine learning, Plink, and PRSice, respectively. It also shows the count of phenotypes for which this combination of tool and hyperparameter generated the best AUC. Each cell reports AUC in percent for a phenotype. Cells for which the value is 0 mean that the tool and hyperparameter combination did not yield the best AUC.

Among ML algorithms, XGBoost is the best algorithm, which produces the best results for 11 phenotypes. It works by combining multiple weak models (Decision trees) to create a strong predictive model. XGBoost has been used for genotype-phenotype prediction [98, 74].

Among DL algorithms, ANN, BILSTM and BILSTM-GRU produced the best results for 4, 3 and 3 phenotypes. ANN and recurrent neural networks have been extensively used for genotype phenotype prediction [99, 61, 100, 101]. SNPs are passed to the ANN model, which finds the non-linear relationship among SNPs to improve the classification score. In contrast, recurrent neural networks capture the changes in the SNPs sequence for classification. BILSTM creates two instances of LSTM, and SNPs are processed in the reverse order for the second instance. This combination of two instances can assist in producing a better performance for datasets with limited samples.

Among PRS tools, Plink is the best tool, producing the best results for 25 phenotypes. Plink produces PRS by summing the product of each SNP’s genotype data and effect size. We noticed that Plink worked well when the clumping threshold was 0.1, which means that it included SNPs in weaker LD, capturing more potentially correlated variants. Lassosum worked well for mutations in pruning parameters, as pruning reduces the number of variants and focuses on those that are more informative or relevant for a particular analysis. This observation aligns with the working principle of the lasso, as it discards SNPs with weights near 0 and considers SNPs relevant to the analysis. PRSice works by summing the product of each SNP’s genotype data and effect size and dividing the sum by the number of SNPs. In our dataset, where the genotype rate is low, PRSice does not account for this missing data. It divides the final sum by the actual number of SNPs, resulting in suboptimal PRS..

Please see supplementary section 2 for detailed results from the ML/DL algorithms and PRS tools.

## Conclusion

openSNP is a growing genetic database publicly available and contains various phenotypes. In this manuscript, we extracted binary phenotypes from the openSNP data and benchmarked the performance of various ML/DL algorithms and PRS tools. We observed that for 36 phenotypes, PRS tools are better; however, for 44 phenotypes, ML/DL tools are better. We discovered that the best-performing machine/deep learning models across all phenotypes were XGBoost and ANN. The incorporation of RNNs in genome imputation justified their inclusion in genotype-phenotype prediction. These models were previously utilized for genotype data prediction, as demonstrated in studies by Pouladi et al. [89] and Srinivasu et al. [90] Notably, RNN networks, particularly stack models, yielded positive results for specific phenotypes, suggesting their suitability for genotype-phenotype prediction. However, it is essential to note that these models may lack direct biological interpretability. The best-performing model depends on several factors like genotype data, quality control steps, phenotype value transformation or labeling, phenotype under consideration, clumping and pruning parameters, and the algorithm used for classification, making genotype-phenotype a complex problem, and providing an opportunity to explore other state-of-the-art methods for improvement. openSNP quantity is relatively low and restricts interpretation and reliance on the results. Though, the analysis performed in this manuscript assists in phenotypes and populations for which the genotype data is limited. Transfer learning has been employed to use knowledge from the well-studied population for genotype-phenotype prediction of the under-studied population with limited data available. The methods presented in this paper used brute-force techniques to explore several methods and hyperparameters that may work well given the limited genotype data, making it suitable for the population for which genotype data is limited. We also reported the best model and corresponding hyperparameter, which generated the best AUC for a specific phenotype. For genotype-phenotype prediction, we recommend that the researchers use ANN with the architecture with five layers, as multiple studies suggest. Various combinations of layers and number of neurons can also be tried for pilot testing. The number of SNPs depends on the complexity of the phenotype. Phenotypes with complex genotype architecture usually require SNPs in tens of thousands for training to produce a better performance. Second, among PRS tools, Plink with default clumping and pruning parameters should be initially tried for genotype-phenotype prediction.

There are a few limitations of this study. Clumping, pruning, minor allele frequency, sex discrepancy, SNP-level missingness, Hardy–Weinberg (dis)equilibrium, and linkage disequilibrium were integral parts of the GWAS file construction process [102, 10]. However, lack of gender and population information are inherent limitations of open-source data (openSNP) because the genetic data for such sources stem from Direct-To-Consumer genetic testing (DTC-GT) offered by companies like 23andMe, Family Tree DNA, or Ancestry.com. In openSNP, the majority of respondents (60.33%) came from the USA, followed by Canada (5.17%) and the UK (4.61%). Population stratification is essential for the analysis because allele frequencies can differ between subpopulations, potentially leading to false positive associations and masking true associations. This is especially critical when identifying the actual SNPs associated with diseases. In our case (case-control classification), the presence of false associations degrades the model’s performance. This is evident because, for some phenotypes, the optimal number of SNPs for classification was found to be between 1000 to 10000. This suggests that the SNPs selected by low p-value thresholds may not fully describe the entire phenotype variation. A PRS model can also be entirely based on genetic data, but in our analysis, we included six principal components and gender [103]. Age and other confounding factors were not present in openSNP data.

We planned to improve the performance by constructing a multi-model using data from DL, ML, and PRS tools. The code and analysis presented in this study allow other researchers working on this dataset to reproduce the result and get more insights into the genotype-phenotype association. The associated code for this manuscript is available on https://github.com/MuhammadMuneeb007/Benchmarking-80-openSNP-phenotypes-using-deep-learning-algorithms-and-polygenic-risk-scores-tools. We documented the procedures for benchmarking so that when the dataset grows large in quantity, researchers could use the code associated with this research work to reperform the existing analysis. We used Python, R, and ML/DL libraries, including scikit-learn [81], TensorFlow [104], and Keras [105] for implementing our analysis.

### Key Points

- This study compares the performance of 29 ML algorithms, 80 DL algorithms and hyperparameter combinations, and three PRS tools (675 clumping and pruning parameters) across 80 phenotypes.
- We reported the best-performing algorithm and corresponding hyperparameters for each phenotype. We noticed the XGBoost (machine learning), ANN (deep learning), and Plink (PRS tools) yielded the best performance for most of the phenotypes.
- Despite the limited quantity of data, some algorithms produced quality results suggesting that this analysis can be used for under-studied populations with limited data assisting in precision medicine.
- The optimal performance of ML/DL algorithms and PRS for phenotypes depend on multiple factors, including hyperparameters, algorithms, phenotype under consideration, and data quality and quantity. Why a specific algorithm yields optimal results for a particular phenotype is yet to discover.

## Author contributions statement

M.M. wrote the first draft of the manuscript. M.M. and Y.M. analyzed the results. D.A., S.F., Y.M., and A.H. reviewed and edited the manuscript. All authors contributed to the methodology of the manuscript.

## Data availability

The dataset we used for this study is available from openSNP (https://opensnp.org/) and the code is available on the GitHub repository (https://github.com/MuhammadMuneeb007/Benchmarking-80-openSNP-phenotypes-using-deep-learning-algorithms-and-polygenic-risk-scores-tools).

## Acknowledgments

Not applicable

**Abstract**

Supplementary Material 1.

This document contains details of dataset pre-processing, including cleaning phenotypes values and converting genotype files. It also contains detailed results achieved by each algorithm.

## Dataset

In openSNP, the majority of respondents (60.33%) came from the USA, followed by Canada (5.17%) and the UK (4.61%). An inherent limitation of open-source data (openSNP) is the absence of gender and population information, as this genetic data originates from Direct-To-Consumer genetic testing (DTC-GT) offered by companies like 23andMe, Family Tree DNA, or Ancestry.com. Despite some data limitations, there are compelling reasons to assess this dataset. The openSNP database comprises diverse data from individuals, and its growth is expected to continue in the future [2, 4], necessitating performance evaluation. Furthermore, the dataset offers an abundance of phenotype data, enabling a wide range of analyses. Notably, our study represents one of the first comprehensive and systematic research efforts benchmarking the openSNP data using machine/deep learning and PRS tools. While the dataset may be relatively small in size, it facilitates efficient analyses. As an illustration, we employed 675 PRS predictors for a single phenotype.

### Dataset pre-processing

We considered binary phenotypes from the openSNP dataset [1]. The dataset contains genotype files in 23andMe, DecodeMe, and FamilyTreeDNA file format and phenotype file in which each column represents the phenotype and each row corresponds to a specific person. There are 6,401 genotype files with 668 phenotypes. However, phenotype values for most people are missing, and we discard phenotypes with missing values. The first step was to define two unique values for each phenotype and the number of people for which a particular phenotype exists. The information is available on GitHub (Analysis1.pdf). The next step was to clean the dataset manually. We considered only binary phenotypes and mapped each value for each phenotype to three values: Case, Control, and Unknown.

### Phenotype values transformation

Phenotype label responses by participants are inconsistent, requiring phenotype data cleaning. For instance, inconsistent terminology is used for Right-handedness: “right-handed”, “right”, “Right”, and “R”. We unified these instances to a single phenotype value, so cases and controls can be generated for each phenotype. This process is referred to as phenotype cleaning or transformation [5], and it is a manual and time-consuming process. Along with examples, some other ambiguities in phenotype values are illustrated on GitHub (AmbiguousPhenotypes.pdf).

We mapped phenotypes values to unique instances in two steps to cope with this issue. The first step was to generate the `preTransform file`, which lists the phenotype, unique values for each phenotype, and the number of values for each phenotype. This file is available on GitHub (preTransform.csv) and helps trace the phenotype file’s ambiguities.

The next step was to generate the `postTransform file`, which maps multiple values for each phenotype and phenotype’s values to one of the following values for binary classification. Manual mapping phenotype values may take approximately 3 to 5 days.

- Cases: Yes (Presence of a particular phenotype value).
- Controls: No (Absence of a particular phenotype value).
- U: Unsure, means we are unsure whether people who selected this phenotype fall in Class1 or Class2.
- x: means this phenotype is removed from further analysis because its values contain too many instances or a high difference between the number of cases and control.

The final postTransformed is available on this GitHub (postTransformed.csv).

### Extract samples for each phenotype

Once we had the postTransform file, the next step was to change the values from the preTransform file to the postTransform file in the phenotype file, count the number of cases/controls for each phenotype, and extract genetic files of people for which the corresponding phenotype value is present. The phenotype name, the number of samples of class 1 and class 2, the total number of samples for each phenotype, and the actual class name are available on GitHub (Analysis2.pdf). There are 101 phenotypes we considered for further analysis. The next step is to extract the genetic files for each phenotype and place them in a specific directory for further processing.

### Genetic file format conversion

The genetic files were in 23andme and AncestryDNA file format. We converted all AncestryDNA files to 23andme file format and then converted 23andme file format to Plink file format (bed, bim, fam) [3]. Table 1 shows the phenotypes and the number of samples for each phenotype.

**Table 1:**
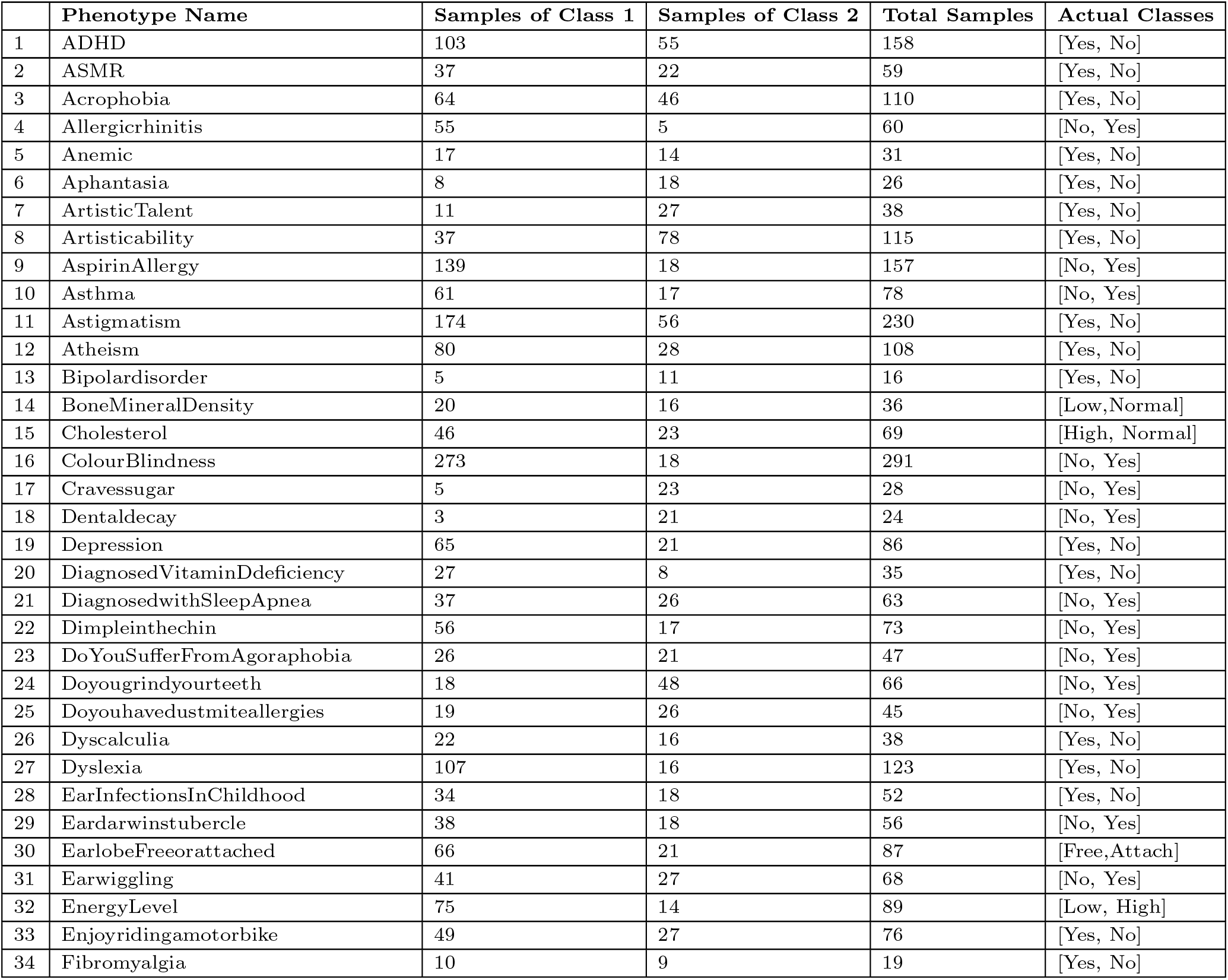
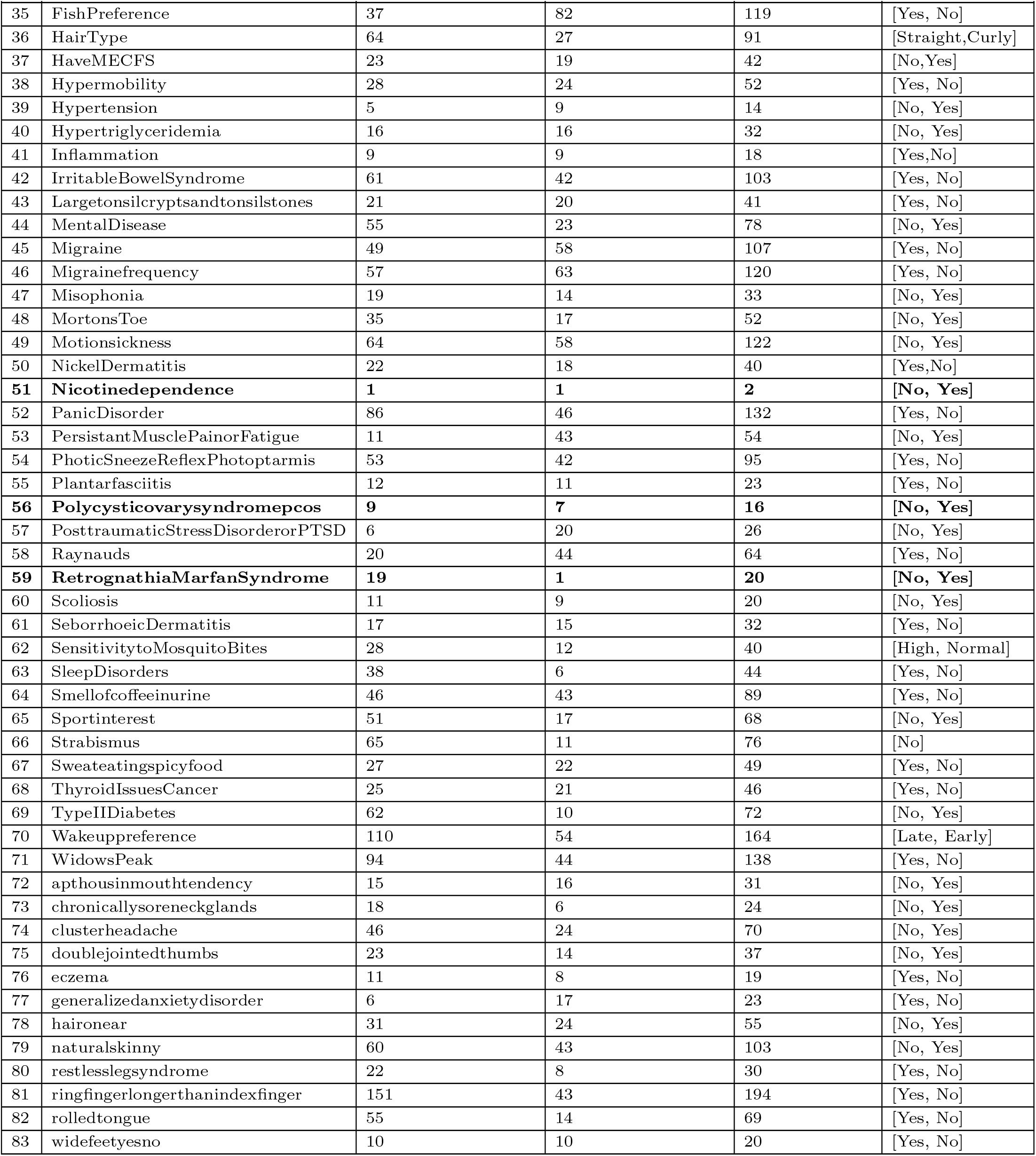
This table shows the phenotypes and the number of samples for each phenotype. The first column reports the phenotype, the second column shows samples from class 1, the third column shows the samples from class 2, the fourth column shows the total number of samples, and the last column shows the actual phenotype values. The bold rows show phenotypes removed by Plink from further analysis due to the low sample size.

## Results

This section contains detailed results from all algorithms. Tables 2, 3, 4, 5 and 6 show the results from machine learning, deep learning, Plink, PRSice, and Lassosum. Table 7 combines the AUC from all tools. Table 8 shows the final results, including the best model and hyperparameters generating the best results.

**Table 2:**
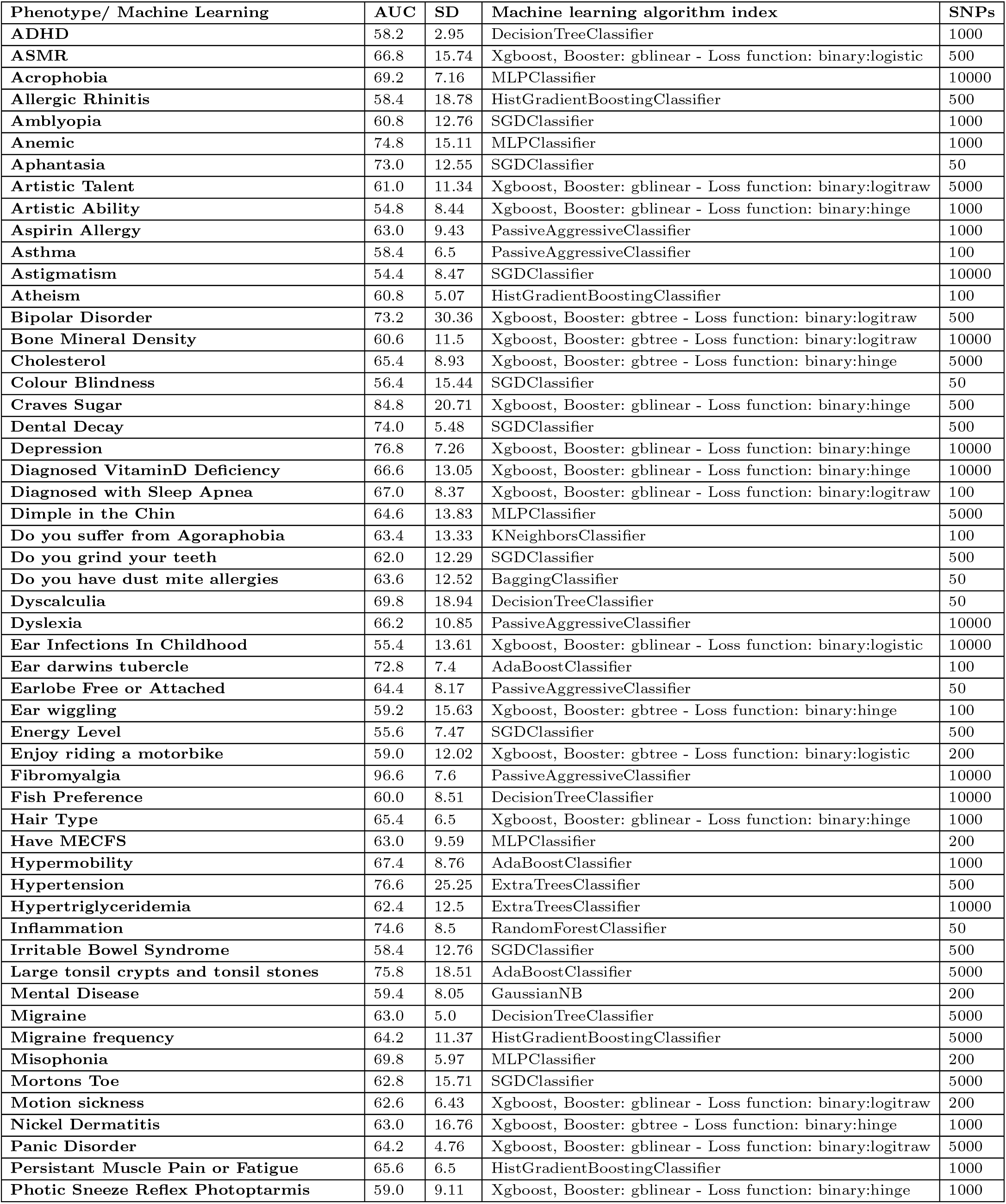
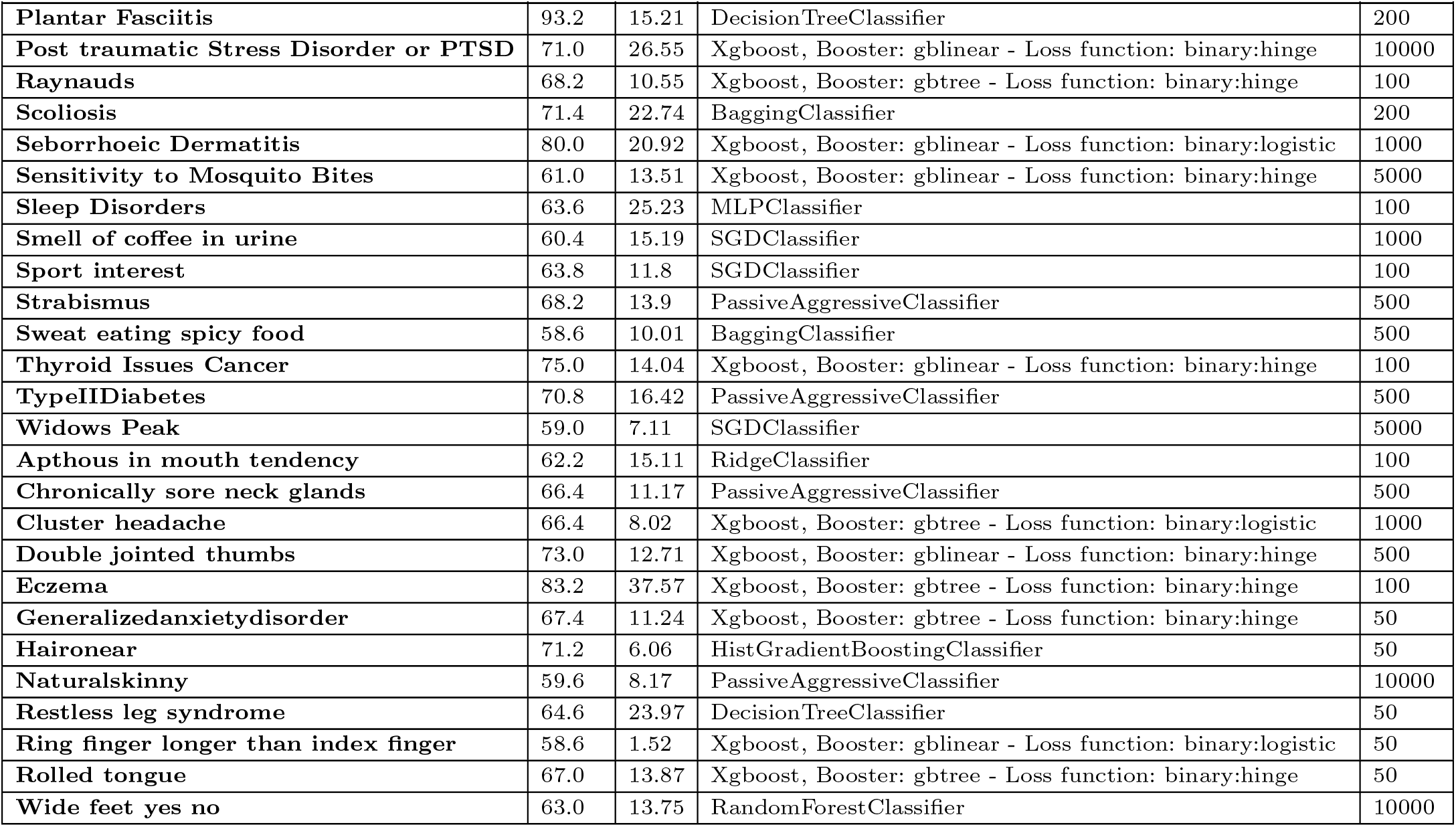
This table shows the results of the machine learning algorithm. The first column reports the phenotype, the second column shows the best AUC, the third column shows the standard deviation and the fourth column shows the model and hyperparameters that generated the best AUC among all other models and hyperparameters. The final column shows the number of SNPs for achieving the best AUC.

| Phenotype/ Machine Learning | AUC | SD | Machine learning algorithm index | SNPs |
| --- | --- | --- | --- | --- |
| ADHD | 58.2 | 2.95 | DecisionTreeClassifier | 1000 |
| ASMR | 66.8 | 15.74 | Xgboost, Booster: gblinear - Loss function: binary:logistic | 500 |
| Acrophobia | 69.2 | 7.16 | MLPClassifier | 10000 |
| Allergic Rhinitis | 58.4 | 18.78 | HistGradientBoostingClassifier | 500 |
| Amblyopia | 60.8 | 12.76 | SGDClassifier | 1000 |
| Anemic | 74.8 | 15.11 | MLPClassifier | 1000 |
| Aphantasia | 73.0 | 12.55 | SGDClassifier | 50 |
| Artistic Talent | 61.0 | 11.34 | Xgboost, Booster: gblinear - Loss function: binary:logitraw | 5000 |
| Artistic Ability | 54.8 | 8.44 | Xgboost, Booster: gblinear - Loss function: binary:hinge | 1000 |
| Aspirin Allergy | 63.0 | 9.43 | PassiveAggressiveClassifier | 1000 |
| Asthma | 58.4 | 6.5 | PassiveAggressiveClassifier | 100 |
| Astigmatism | 54.4 | 8.47 | SGDClassifier | 10000 |
| Atheism | 60.8 | 5.07 | HistGradientBoostingClassifier | 100 |
| Bipolar Disorder | 73.2 | 30.36 | Xgboost, Booster: gbtree - Loss function: binary:logitraw | 500 |
| Bone Mineral Density | 60.6 | 11.5 | Xgboost, Booster: gbtree - Loss function: binary:logitraw | 10000 |
| Cholesterol | 65.4 | 8.93 | Xgboost, Booster: gbtree - Loss function: binary:hinge | 5000 |
| Colour Blindness | 56.4 | 15.44 | SGDClassifier | 50 |
| Craves Sugar | 84.8 | 20.71 | Xgboost, Booster: gblinear - Loss function: binary:hinge | 500 |
| Dental Decay | 74.0 | 5.48 | SGDClassifier | 500 |
| Depression | 76.8 | 7.26 | Xgboost, Booster: gblinear - Loss function: binary:hinge | 10000 |
| Diagnosed VitaminD Deficiency | 66.6 | 13.05 | Xgboost, Booster: gblinear - Loss function: binary:hinge | 10000 |
| Diagnosed with Sleep Apnea | 67.0 | 8.37 | Xgboost, Booster: gblinear - Loss function: binary:logitraw | 100 |
| Dimple in the Chin | 64.6 | 13.83 | MLPClassifier | 5000 |
| Do you suffer from Agoraphobia | 63.4 | 13.33 | KNeighborsClassifier | 100 |
| Do you grind your teeth | 62.0 | 12.29 | SGDClassifier | 500 |
| Do you have dust mite allergies | 63.6 | 12.52 | BaggingClassifier | 50 |
| Dyscalculia | 69.8 | 18.94 | DecisionTreeClassifier | 50 |
| Dyslexia | 66.2 | 10.85 | PassiveAggressiveClassifier | 10000 |
| Ear Infections In Childhood | 55.4 | 13.61 | Xgboost, Booster: gblinear - Loss function: binary:logistic | 10000 |
| Ear darwins tubercle | 72.8 | 7.4 | AdaBoostClassifier | 100 |
| Earlobe Free or Attached | 64.4 | 8.17 | PassiveAggressiveClassifier | 50 |
| Ear wiggling | 59.2 | 15.63 | Xgboost, Booster: gbtree - Loss function: binary:hinge | 100 |
| Energy Level | 55.6 | 7.47 | SGDClassifier | 500 |
| Enjoy riding a motorbike | 59.0 | 12.02 | Xgboost, Booster: gbtree - Loss function: binary:logistic | 200 |
| Fibromyalgia | 96.6 | 7.6 | PassiveAggressiveClassifier | 10000 |
| Fish Preference | 60.0 | 8.51 | DecisionTreeClassifier | 10000 |
| Hair Type | 65.4 | 6.5 | Xgboost, Booster: gblinear - Loss function: binary:hinge | 1000 |
| Have MECFS | 63.0 | 9.59 | MLPClassifier | 200 |
| Hypermobility | 67.4 | 8.76 | AdaBoostClassifier | 1000 |
| Hypertension | 76.6 | 25.25 | ExtraTreesClassifier | 500 |
| Hypertriglyceridemia | 62.4 | 12.5 | ExtraTreesClassifier | 10000 |
| Inflammation | 74.6 | 8.5 | RandomForestClassifier | 50 |
| Irritable Bowel Syndrome | 58.4 | 12.76 | SGDClassifier | 500 |
| Large tonsil crypts and tonsil stones | 75.8 | 18.51 | AdaBoostClassifier | 5000 |
| Mental Disease | 59.4 | 8.05 | GaussianNB | 200 |
| Migraine | 63.0 | 5.0 | DecisionTreeClassifier | 5000 |
| Migraine frequency | 64.2 | 11.37 | HistGradientBoostingClassifier | 5000 |
| Misophonia | 69.8 | 5.97 | MLPClassifier | 200 |
| Mortons Toe | 62.8 | 15.71 | SGDClassifier | 5000 |
| Motion sickness | 62.6 | 6.43 | Xgboost, Booster: gblinear - Loss function: binary:logitraw | 200 |
| Nickel Dermatitis | 63.0 | 16.76 | Xgboost, Booster: gbtree - Loss function: binary:hinge | 1000 |
| Panic Disorder | 64.2 | 4.76 | Xgboost, Booster: gblinear - Loss function: binary:logitraw | 5000 |
| Persistent Muscle Pain or Fatigue | 65.6 | 6.5 | HistGradientBoostingClassifier | 1000 |
| Photic Sneeze Reflex Photoptarmis | 59.0 | 9.11 | Xgboost, Booster: gblinear - Loss function: binary:hinge | 1000 |
| Plantar Fasciitis | 93.2 | 15.21 | DecisionTreeClassifier | 200 |
| Post traumatic Stress Disorder or PTSD | 71.0 | 26.55 | Xgboost, Booster: gblinear - Loss function: binary:hinge | 10000 |
| Raynauds | 68.2 | 10.55 | Xgboost, Booster: gbtree - Loss function: binary:hinge | 100 |
| Scoliosis | 71.4 | 22.74 | BaggingClassifier | 200 |
| Seborrhoeic Dermatitis | 80.0 | 20.92 | Xgboost, Booster: gblinear - Loss function: binary:logistic | 1000 |
| Sensitivity to Mosquito Bites | 61.0 | 13.51 | Xgboost, Booster: gblinear - Loss function: binary:hinge | 5000 |
| Sleep Disorders | 63.6 | 25.23 | MLPClassifier | 100 |
| Smell of coffee in urine | 60.4 | 15.19 | SGDClassifier | 1000 |
| Sport interest | 63.8 | 11.8 | SGDClassifier | 100 |
| Strabismus | 68.2 | 13.9 | PassiveAggressiveClassifier | 500 |
| Sweat eating spicy food | 58.6 | 10.01 | BaggingClassifier | 500 |
| Thyroid Issues Cancer | 75.0 | 14.04 | Xgboost, Booster: gblinear - Loss function: binary:hinge | 100 |
| TypeIIDiabetes | 70.8 | 16.42 | PassiveAggressiveClassifier | 500 |
| Widows Peak | 59.0 | 7.11 | SGDClassifier | 5000 |
| Apthous in mouth tendency | 62.2 | 15.11 | RidgeClassifier | 100 |
| Chronically sore neck glands | 66.4 | 11.17 | PassiveAggressiveClassifier | 500 |
| Cluster headache | 66.4 | 8.02 | Xgboost, Booster: gbtree - Loss function: binary:logistic | 1000 |
| Double jointed thumbs | 73.0 | 12.71 | Xgboost, Booster: gblinear - Loss function: binary:hinge | 500 |
| Eczema | 83.2 | 37.57 | Xgboost, Booster: gbtree - Loss function: binary:hinge | 100 |
| Generalizedanxietydisorder | 67.4 | 11.24 | Xgboost, Booster: gbtree - Loss function: binary:hinge | 50 |
| Haironear | 71.2 | 6.06 | HistGradientBoostingClassifier | 50 |
| Naturalskinny | 59.6 | 8.17 | PassiveAggressiveClassifier | 10000 |
| Restless leg syndrome | 64.6 | 23.97 | DecisionTreeClassifier | 50 |
| Ring finger longer than index finger | 58.6 | 1.52 | Xgboost, Booster: gblinear - Loss function: binary:logistic | 50 |
| Rolled tongue | 67.0 | 13.87 | Xgboost, Booster: gbtree - Loss function: binary:hinge | 50 |
| Wide feet yes no | 63.0 | 13.75 | RandomForestClassifier | 10000 |

**Table 3:**
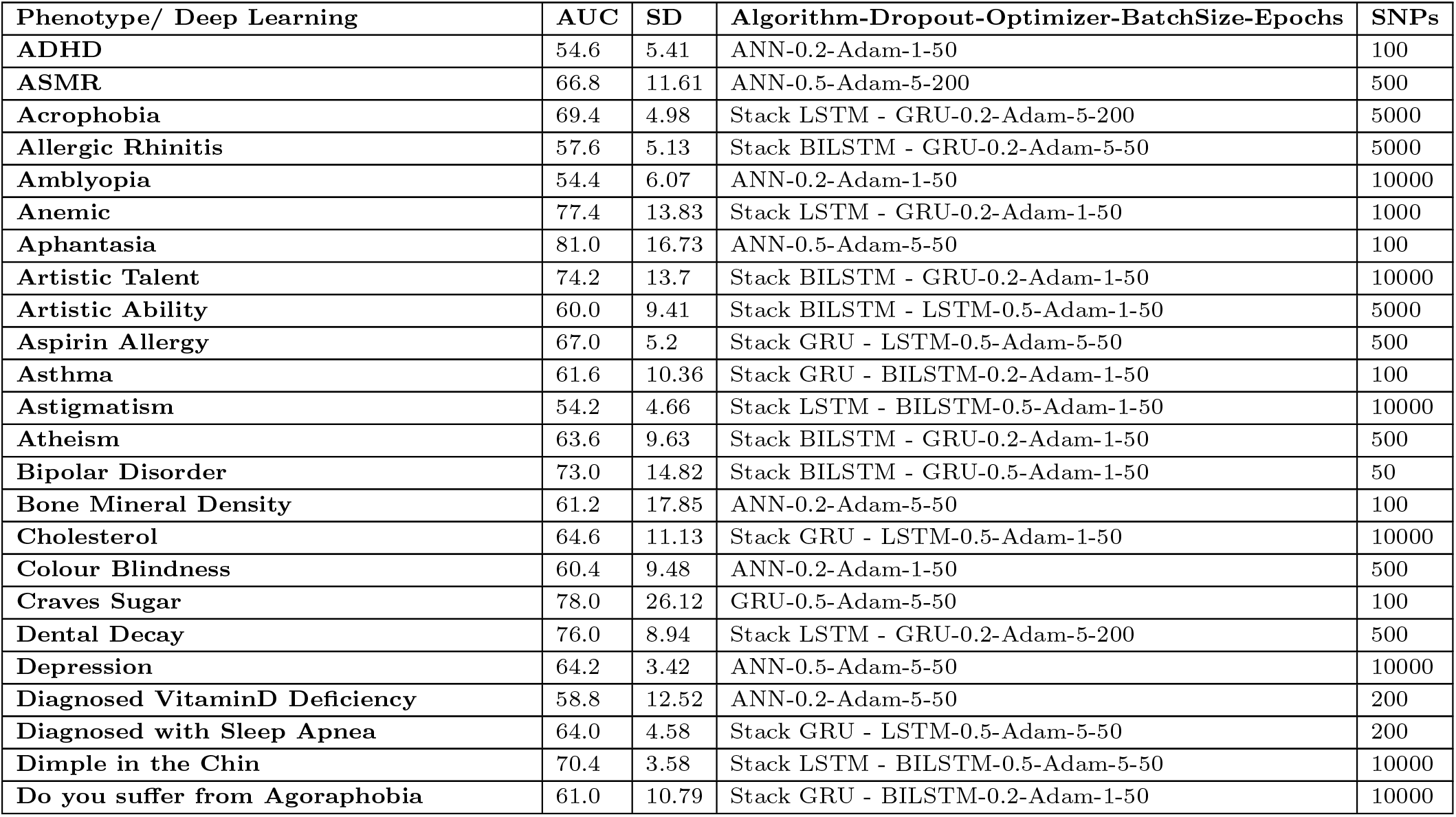
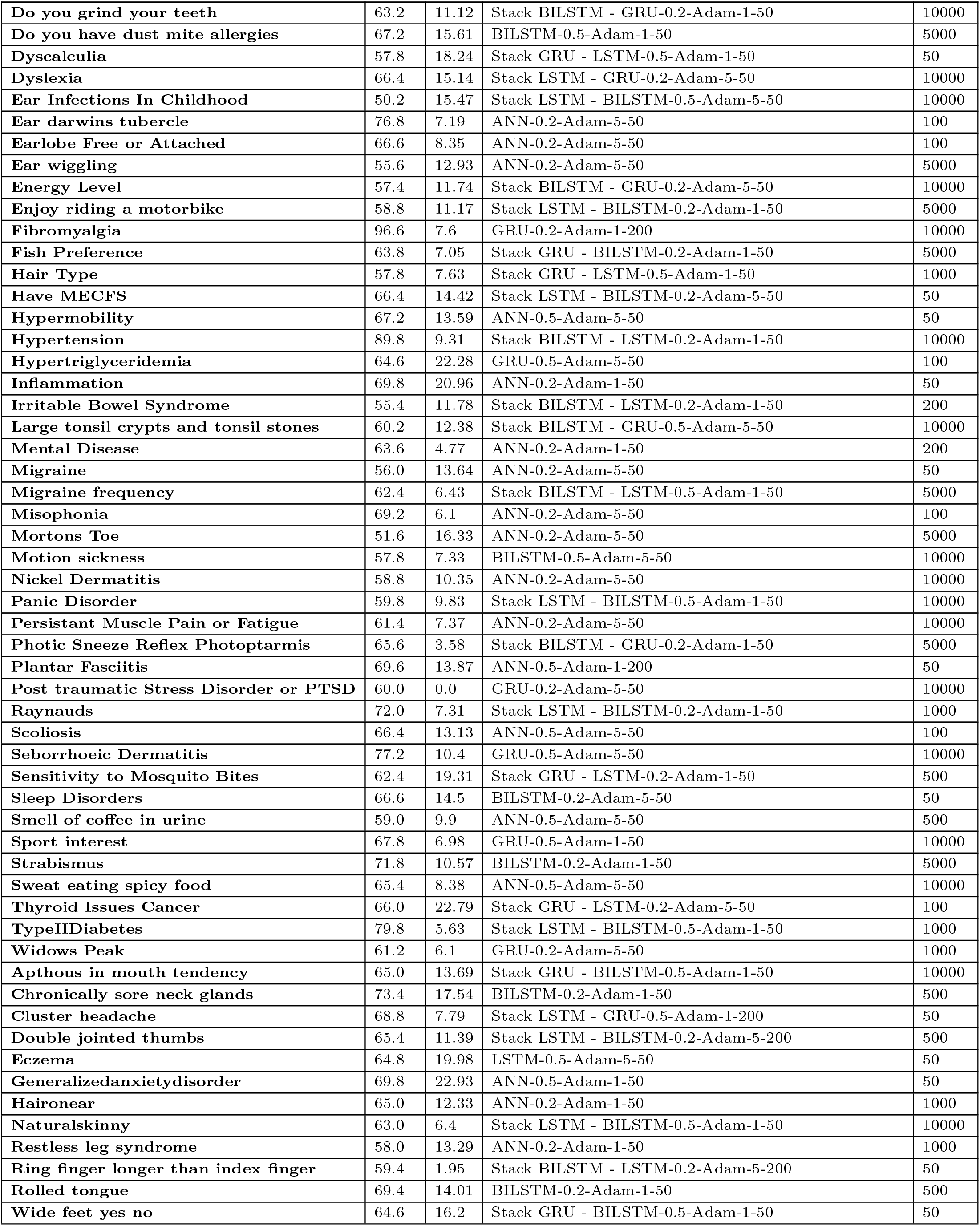
This table shows the results of the deep learning algorithm. The first column reports the phenotype, the second column shows the best AUC, the third column shows the standard deviation and the fourth column shows the model and hyperparameters that generated the best AUC among all other models and hyperparameters. The final column shows the number of SNPs for achieving the best AUC.

| Phenotype/ Deep Learning | AUC | SD | Algorithm-Dropout-Optimizer-BatchSize-Epochs | SNPs |
| --- | --- | --- | --- | --- |
| ADHD | 54.6 | 5.41 | ANN-0.2-Adam-1-50 | 100 |
| ASMR | 66.8 | 11.61 | ANN-0.5-Adam-5-200 | 500 |
| Acrophobia | 69.4 | 4.98 | Stack LSTM - GRU-0.2-Adam-5-200 | 5000 |
| Allergic Rhinitis | 57.6 | 5.13 | Stack BiLSTM - GRU-0.2-Adam-5-50 | 5000 |
| Amblyopia | 54.4 | 6.07 | ANN-0.2-Adam-1-50 | 10000 |
| Anemic | 77.4 | 13.83 | Stack LSTM - GRU-0.2-Adam-1-50 | 1000 |
| Aphantasia | 81.0 | 16.73 | ANN-0.5-Adam-5-50 | 100 |
| Artistic Talent | 74.2 | 13.7 | Stack BiLSTM - GRU-0.2-Adam-1-50 | 10000 |
| Artistic Ability | 60.0 | 9.41 | Stack BiLSTM - LSTM-0.5-Adam-1-50 | 5000 |
| Aspirin Allergy | 67.0 | 5.2 | Stack GRU - LSTM-0.5-Adam-5-50 | 500 |
| Asthma | 61.6 | 10.36 | Stack GRU - BiLSTM-0.2-Adam-1-50 | 100 |
| Astigmatism | 54.2 | 4.66 | Stack LSTM - BiLSTM-0.5-Adam-1-50 | 10000 |
| Atheism | 63.6 | 9.63 | Stack BiLSTM - GRU-0.2-Adam-1-50 | 500 |
| Bipolar Disorder | 73.0 | 14.82 | Stack BiLSTM - GRU-0.5-Adam-1-50 | 50 |
| Bone Mineral Density | 61.2 | 17.85 | ANN-0.2-Adam-5-50 | 100 |
| Cholesterol | 64.6 | 11.13 | Stack GRU - LSTM-0.5-Adam-1-50 | 10000 |
| Colour Blindness | 60.4 | 9.48 | ANN-0.2-Adam-1-50 | 500 |
| Craves Sugar | 78.0 | 26.12 | GRU-0.5-Adam-5-50 | 100 |
| Dental Decay | 76.0 | 8.94 | Stack LSTM - GRU-0.2-Adam-5-200 | 500 |
| Depression | 64.2 | 3.42 | ANN-0.5-Adam-5-50 | 10000 |
| Diagnosed VitaminD Deficiency | 58.8 | 12.52 | ANN-0.2-Adam-5-50 | 200 |
| Diagnosed with Sleep Apnea | 64.0 | 4.58 | Stack GRU - LSTM-0.5-Adam-5-50 | 200 |
| Dimple in the Chin | 70.4 | 3.58 | Stack LSTM - BiLSTM-0.5-Adam-5-50 | 10000 |
| Do you suffer from Agoraphobia | 61.0 | 10.79 | Stack GRU - BiLSTM-0.2-Adam-1-50 | 10000 |
| Do you grind your teeth | 63.2 | 11.12 | Stack BILSTM - GRU-0.2-Adam-1-50 | 10000 |
| Do you have dust mite allergies | 67.2 | 15.61 | BILSTM-0.5-Adam-1-50 | 5000 |
| Dyscalculia | 57.8 | 18.24 | Stack GRU - LSTM-0.5-Adam-1-50 | 50 |
| Dyslexia | 66.4 | 15.14 | Stack LSTM - GRU-0.2-Adam-5-50 | 10000 |
| Ear Infections In Childhood | 50.2 | 15.47 | Stack LSTM - BILSTM-0.5-Adam-5-50 | 10000 |
| Ear darwins tubercle | 76.8 | 7.19 | ANN-0.2-Adam-5-50 | 100 |
| Earlobe Free or Attached | 66.6 | 8.35 | ANN-0.2-Adam-5-50 | 100 |
| Ear wiggling | 55.6 | 12.93 | ANN-0.2-Adam-5-50 | 5000 |
| Energy Level | 57.4 | 11.74 | Stack BILSTM - GRU-0.2-Adam-5-50 | 10000 |
| Enjoy riding a motorbike | 58.8 | 11.17 | Stack LSTM - BILSTM-0.2-Adam-1-50 | 5000 |
| Fibromyalgia | 96.6 | 7.6 | GRU-0.2-Adam-1-200 | 10000 |
| Fish Preference | 63.8 | 7.05 | Stack GRU - BILSTM-0.2-Adam-1-50 | 5000 |
| Hair Type | 57.8 | 7.63 | Stack GRU - LSTM-0.5-Adam-1-50 | 1000 |
| Have MECFS | 66.4 | 14.42 | Stack LSTM - BILSTM-0.2-Adam-5-50 | 50 |
| Hypermobility | 67.2 | 13.59 | ANN-0.5-Adam-5-50 | 50 |
| Hypertension | 89.8 | 9.31 | Stack BILSTM - LSTM-0.2-Adam-1-50 | 10000 |
| Hypertriglyceridemia | 64.6 | 22.28 | GRU-0.5-Adam-5-50 | 100 |
| Inflammation | 69.8 | 20.96 | ANN-0.2-Adam-1-50 | 50 |
| Irritable Bowel Syndrome | 55.4 | 11.78 | Stack BILSTM - LSTM-0.2-Adam-1-50 | 200 |
| Large tonsil crypts and tonsil stones | 60.2 | 12.38 | Stack BILSTM - GRU-0.5-Adam-5-50 | 10000 |
| Mental Disease | 63.6 | 4.77 | ANN-0.2-Adam-1-50 | 200 |
| Migraine | 56.0 | 13.64 | ANN-0.2-Adam-5-50 | 50 |
| Migraine frequency | 62.4 | 6.43 | Stack BILSTM - LSTM-0.5-Adam-1-50 | 5000 |
| Misophonia | 69.2 | 6.1 | ANN-0.2-Adam-5-50 | 100 |
| Mortons Toe | 51.6 | 16.33 | ANN-0.2-Adam-5-50 | 5000 |
| Motion sickness | 57.8 | 7.33 | BILSTM-0.5-Adam-5-50 | 10000 |
| Nickel Dermatitis | 58.8 | 10.35 | ANN-0.2-Adam-5-50 | 10000 |
| Panic Disorder | 59.8 | 9.83 | Stack LSTM - BILSTM-0.5-Adam-1-50 | 10000 |
| Persistent Muscle Pain or Fatigue | 61.4 | 7.37 | ANN-0.2-Adam-5-50 | 10000 |
| Photoc Sneeze Reflex Photoptarmis | 65.6 | 3.58 | Stack BILSTM - GRU-0.2-Adam-1-50 | 5000 |
| Plantar Fasciitis | 69.6 | 13.87 | ANN-0.5-Adam-1-200 | 50 |
| Post traumatic Stress Disorder or PTSD | 60.0 | 0.0 | GRU-0.2-Adam-5-50 | 10000 |
| Raynauds | 72.0 | 7.31 | Stack LSTM - BILSTM-0.2-Adam-1-50 | 1000 |
| Scoliosis | 66.4 | 13.13 | ANN-0.5-Adam-5-50 | 100 |
| Seborrhoeic Dermatitis | 77.2 | 10.4 | GRU-0.5-Adam-5-50 | 10000 |
| Sensitivity to Mosquito Bites | 62.4 | 19.31 | Stack GRU - LSTM-0.2-Adam-1-50 | 500 |
| Sleep Disorders | 66.6 | 14.5 | BILSTM-0.2-Adam-5-50 | 50 |
| Smell of coffee in urine | 59.0 | 9.9 | ANN-0.5-Adam-5-50 | 500 |
| Sport interest | 67.8 | 6.98 | GRU-0.5-Adam-1-50 | 10000 |
| Strabismus | 71.8 | 10.57 | BILSTM-0.2-Adam-1-50 | 5000 |
| Sweat eating spicy food | 65.4 | 8.38 | ANN-0.5-Adam-5-50 | 10000 |
| Thyroid Issues Cancer | 66.0 | 22.79 | Stack GRU - LSTM-0.2-Adam-5-50 | 100 |
| TypeIIDiabetes | 79.8 | 5.63 | Stack LSTM - BILSTM-0.5-Adam-1-50 | 1000 |
| Widows Peak | 61.2 | 6.1 | GRU-0.2-Adam-5-50 | 1000 |
| Apthous in mouth tendency | 65.0 | 13.69 | Stack GRU - BILSTM-0.5-Adam-1-50 | 10000 |
| Chronically sore neck glands | 73.4 | 17.54 | BILSTM-0.2-Adam-1-50 | 500 |
| Cluster headache | 68.8 | 7.79 | Stack LSTM - GRU-0.5-Adam-1-200 | 50 |
| Double jointed thumbs | 65.4 | 11.39 | Stack LSTM - BILSTM-0.2-Adam-5-200 | 500 |
| Eczema | 64.8 | 19.98 | LSTM-0.5-Adam-5-50 | 50 |
| Generalizedanxietydisorder | 69.8 | 22.93 | ANN-0.5-Adam-1-50 | 50 |
| Hairnear | 65.0 | 12.33 | ANN-0.2-Adam-1-50 | 1000 |
| Natural skinny | 63.0 | 6.4 | Stack LSTM - BILSTM-0.5-Adam-1-50 | 10000 |
| Restless leg syndrome | 58.0 | 13.29 | ANN-0.2-Adam-1-50 | 1000 |
| Ring finger longer than index finger | 59.4 | 1.95 | Stack BILSTM - LSTM-0.2-Adam-5-200 | 50 |
| Rolled tongue | 69.4 | 14.01 | BILSTM-0.2-Adam-1-50 | 500 |
| Wide feet yes no | 64.6 | 16.2 | Stack GRU - BILSTM-0.5-Adam-1-50 | 50 |

| Phenotype/ Deep Learning | AUC | SD | Algorithm-Dropout-Optimizer-BatchSize-Epochs | SNPs |
| --- | --- | --- | --- | --- |

**Table 4:**
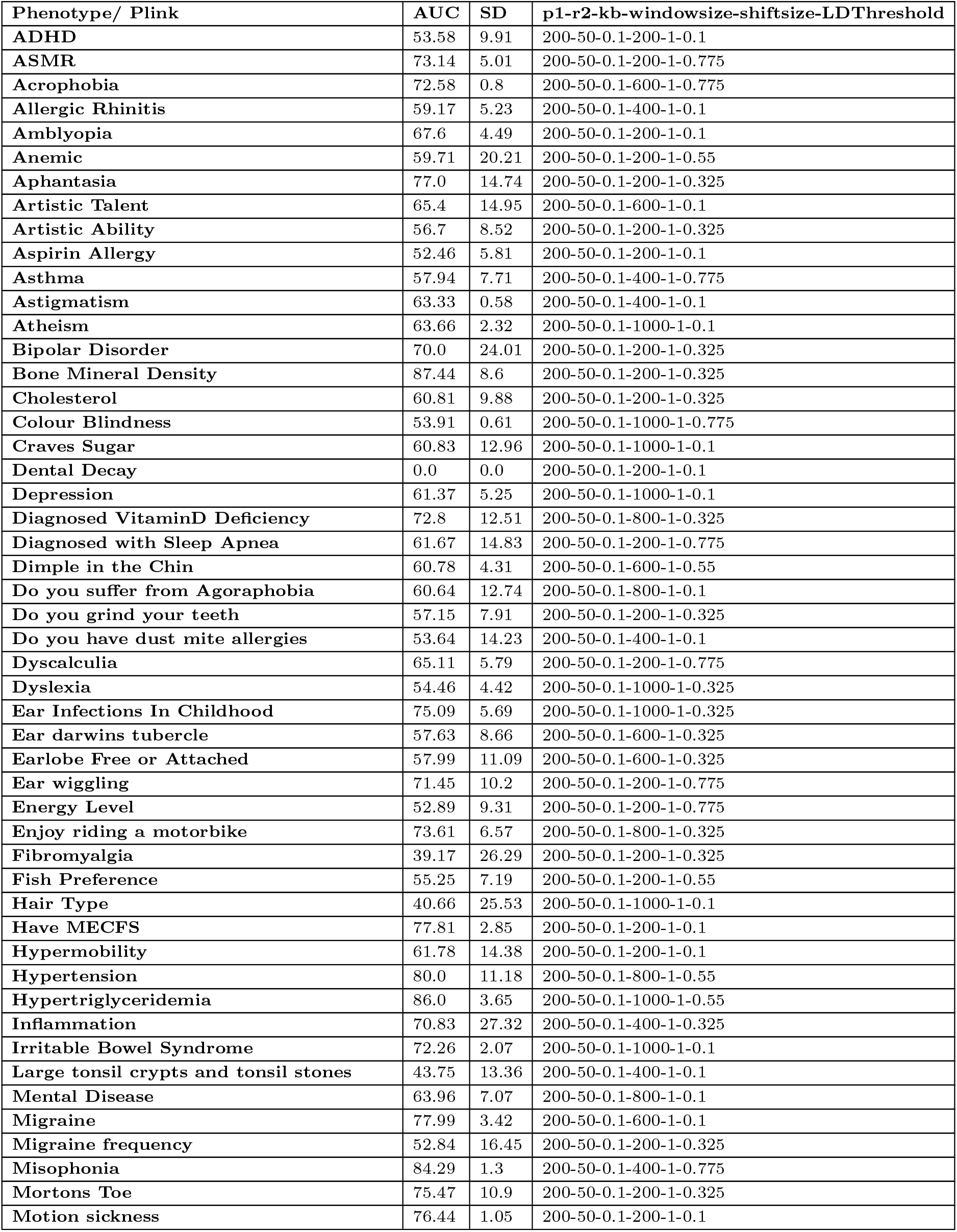
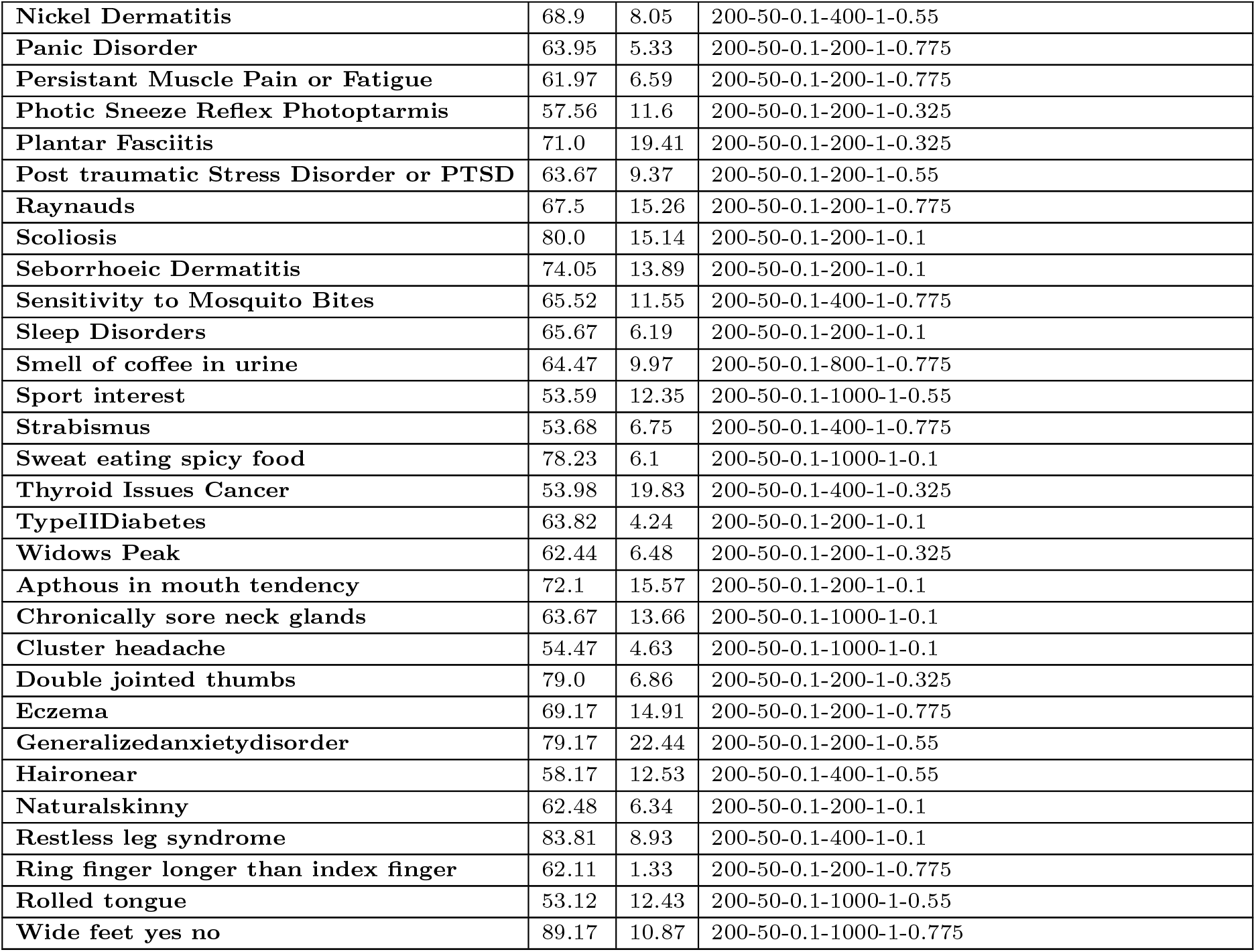
This table shows the Plink results. The first column reports the phenotype, the second column shows the best AUC, the third column shows the standard deviation and the fourth column shows the hyperparameters or predictors that generated the best AUC among all other models and hyperparameters.

| Phenotype/ Plink | AUC | SD | p1-r2-kb-window-size-shift-size-LDThreshold |
| --- | --- | --- | --- |
| ADHD | 53.58 | 9.91 | 200-50-0.1-200-1-0.1 |
| ASMR | 73.14 | 5.01 | 200-50-0.1-200-1-0.775 |
| Acrophobia | 72.58 | 0.8 | 200-50-0.1-600-1-0.775 |
| Allergic Rhinitis | 59.17 | 5.23 | 200-50-0.1-400-1-0.1 |
| Amblyopia | 67.6 | 4.49 | 200-50-0.1-200-1-0.1 |
| Anemic | 59.71 | 20.21 | 200-50-0.1-200-1-0.55 |
| Aphantasia | 77.0 | 14.74 | 200-50-0.1-200-1-0.325 |
| Artistic Talent | 65.4 | 14.95 | 200-50-0.1-600-1-0.1 |
| Artistic Ability | 56.7 | 8.52 | 200-50-0.1-200-1-0.325 |
| Aspirin Allergy | 52.46 | 5.81 | 200-50-0.1-200-1-0.1 |
| Asthma | 57.94 | 7.71 | 200-50-0.1-400-1-0.775 |
| Astigmatism | 63.33 | 0.58 | 200-50-0.1-400-1-0.1 |
| Atheism | 63.66 | 2.32 | 200-50-0.1-1000-1-0.1 |
| Bipolar Disorder | 70.0 | 24.01 | 200-50-0.1-200-1-0.325 |
| Bone Mineral Density | 87.44 | 8.6 | 200-50-0.1-200-1-0.325 |
| Cholesterol | 60.81 | 9.88 | 200-50-0.1-200-1-0.325 |
| Colour Blindness | 53.91 | 0.61 | 200-50-0.1-1000-1-0.775 |
| Craves Sugar | 60.83 | 12.96 | 200-50-0.1-1000-1-0.1 |
| Dental Decay | 0.0 | 0.0 | 200-50-0.1-200-1-0.1 |
| Depression | 61.37 | 5.25 | 200-50-0.1-1000-1-0.1 |
| Diagnosed VitaminD Deficiency | 72.8 | 12.51 | 200-50-0.1-800-1-0.325 |
| Diagnosed with Sleep Apnea | 61.67 | 14.83 | 200-50-0.1-200-1-0.775 |
| Dimple in the Chin | 60.78 | 4.31 | 200-50-0.1-600-1-0.55 |
| Do you suffer from Agoraphobia | 60.64 | 12.74 | 200-50-0.1-800-1-0.1 |
| Do you grind your teeth | 57.15 | 7.91 | 200-50-0.1-200-1-0.325 |
| Do you have dust mite allergies | 53.64 | 14.23 | 200-50-0.1-400-1-0.1 |
| Dyscalculia | 65.11 | 5.79 | 200-50-0.1-200-1-0.775 |
| Dyslexia | 54.46 | 4.42 | 200-50-0.1-1000-1-0.325 |
| Ear Infections In Childhood | 75.09 | 5.69 | 200-50-0.1-1000-1-0.325 |
| Ear darwins tubercle | 57.63 | 8.66 | 200-50-0.1-600-1-0.325 |
| Earlobe Free or Attached | 57.99 | 11.09 | 200-50-0.1-600-1-0.325 |
| Ear wiggling | 71.45 | 10.2 | 200-50-0.1-200-1-0.775 |
| Energy Level | 52.89 | 9.31 | 200-50-0.1-200-1-0.775 |
| Enjoy riding a motorbike | 73.61 | 6.57 | 200-50-0.1-800-1-0.325 |
| Fibromyalgia | 39.17 | 26.29 | 200-50-0.1-200-1-0.325 |
| Fish Preference | 55.25 | 7.19 | 200-50-0.1-200-1-0.55 |
| Hair Type | 40.66 | 25.53 | 200-50-0.1-1000-1-0.1 |
| Have MECFS | 77.81 | 2.85 | 200-50-0.1-200-1-0.1 |
| Hypermobility | 61.78 | 14.38 | 200-50-0.1-200-1-0.1 |
| Hypertension | 80.0 | 11.18 | 200-50-0.1-800-1-0.55 |
| Hypertriglyceridemia | 86.0 | 3.65 | 200-50-0.1-1000-1-0.55 |
| Inflammation | 70.83 | 27.32 | 200-50-0.1-400-1-0.325 |
| Irritable Bowel Syndrome | 72.26 | 2.07 | 200-50-0.1-1000-1-0.1 |
| Large tonsil crypts and tonsil stones | 43.75 | 13.36 | 200-50-0.1-400-1-0.1 |
| Mental Disease | 63.96 | 7.07 | 200-50-0.1-800-1-0.1 |
| Migraine | 77.99 | 3.42 | 200-50-0.1-600-1-0.1 |
| Migraine frequency | 52.84 | 16.45 | 200-50-0.1-200-1-0.325 |
| Misophonia | 84.29 | 1.3 | 200-50-0.1-400-1-0.775 |
| Mortons Toe | 75.47 | 10.9 | 200-50-0.1-200-1-0.325 |
| Motion sickness | 76.44 | 1.05 | 200-50-0.1-200-1-0.1 |
| Nickel Dermatitis | 68.9 | 8.05 | 200-50-0.1-400-1-0.55 |
| Panic Disorder | 63.95 | 5.33 | 200-50-0.1-200-1-0.775 |
| Persistent Muscle Pain or Fatigue | 61.97 | 6.59 | 200-50-0.1-200-1-0.775 |
| Photoc Sneeze Reflex Photophobia | 57.56 | 11.6 | 200-50-0.1-200-1-0.325 |
| Plantar Fasciitis | 71.0 | 19.41 | 200-50-0.1-200-1-0.325 |
| Post traumatic Stress Disorder or PTSD | 63.67 | 9.37 | 200-50-0.1-200-1-0.55 |
| Raynauds | 67.5 | 15.26 | 200-50-0.1-200-1-0.775 |
| Scoliosis | 80.0 | 15.14 | 200-50-0.1-200-1-0.1 |
| Seborrheic Dermatitis | 74.05 | 13.89 | 200-50-0.1-200-1-0.1 |
| Sensitivity to Mosquito Bites | 65.52 | 11.55 | 200-50-0.1-400-1-0.775 |
| Sleep Disorders | 65.67 | 6.19 | 200-50-0.1-200-1-0.1 |
| Smell of coffee in urine | 64.47 | 9.97 | 200-50-0.1-800-1-0.775 |
| Sport interest | 53.59 | 12.35 | 200-50-0.1-1000-1-0.55 |
| Strabismus | 53.68 | 6.75 | 200-50-0.1-400-1-0.775 |
| Sweat eating spicy food | 78.23 | 6.1 | 200-50-0.1-1000-1-0.1 |
| Thyroid Issues Cancer | 53.98 | 19.83 | 200-50-0.1-400-1-0.325 |
| TypeII Diabetes | 63.82 | 4.24 | 200-50-0.1-200-1-0.1 |
| Widows Peak | 62.44 | 6.48 | 200-50-0.1-200-1-0.325 |
| Apthous in mouth tendency | 72.1 | 15.57 | 200-50-0.1-200-1-0.1 |
| Chronically sore neck glands | 63.67 | 13.66 | 200-50-0.1-1000-1-0.1 |
| Cluster headache | 54.47 | 4.63 | 200-50-0.1-1000-1-0.1 |
| Double jointed thumbs | 79.0 | 6.86 | 200-50-0.1-200-1-0.325 |
| Eczema | 69.17 | 14.91 | 200-50-0.1-200-1-0.775 |
| Generalized anxiety disorder | 79.17 | 22.44 | 200-50-0.1-200-1-0.55 |
| Hair on ear | 58.17 | 12.53 | 200-50-0.1-400-1-0.55 |
| Natural skinny | 62.48 | 6.34 | 200-50-0.1-200-1-0.1 |
| Restless leg syndrome | 83.81 | 8.93 | 200-50-0.1-400-1-0.1 |
| Ring finger longer than index finger | 62.11 | 1.33 | 200-50-0.1-200-1-0.775 |
| Rolled tongue | 53.12 | 12.43 | 200-50-0.1-1000-1-0.55 |
| Wide feet yes no | 89.17 | 10.87 | 200-50-0.1-1000-1-0.775 |

**Table 5:** This table shows the PRSice results. The first column reports the phenotype, the second column shows the best AUC, the third column shows the standard deviation and the fourth column shows the hyperparameters or predictors that generated the best AUC among all other models and hyperparameters.

| Phenotype/ PRSice | AUC | SD | p1-r2-kb-window-size-shift-size-LDThreshold |
| --- | --- | --- | --- |
| ADHD | 40.38 | 16.77 | 200-50-0.1-200-1-0.1 |
| ASMR | 53.0 | 10.12 | 200-50-0.1-200-1-0.1 |
| Acrophobia | 27.42 | 0.8 | 200-50-0.1-200-1-0.1 |
| Allergic Rhinitis | 55.67 | 1.21 | 200-50-0.1-200-1-0.1 |
| Amblyopia | 64.29 | 5.77 | 200-50-0.1-200-1-0.1 |
| Anemic | 50.0 | 9.43 | 200-50-0.1-200-1-0.1 |
| Aphantasia | 30.33 | 22.59 | 200-50-0.1-200-1-0.1 |
| Artistic Talent | 39.42 | 15.07 | 200-50-0.1-200-1-0.1 |
| Artistic Ability | 41.33 | 12.0 | 200-50-0.1-200-1-0.1 |
| Aspirin Allergy | 45.89 | 23.28 | 200-50-0.1-200-1-0.1 |
| Asthma | 48.72 | 13.78 | 200-50-0.1-200-1-0.1 |
| Astigmatism | 40.98 | 7.91 | 200-50-0.1-200-1-0.1 |
| Atheism | 57.54 | 8.72 | 200-50-0.1-200-1-0.1 |
| Bipolar Disorder | 53.33 | 26.09 | 200-50-0.1-200-1-0.1 |
| Bone Mineral Density | 62.33 | 23.42 | 200-50-0.1-200-1-0.1 |
| Cholesterol | 51.43 | 14.8 | 200-50-0.1-200-1-0.1 |
| Colour Blindness | 52.48 | 5.17 | 200-50-0.1-200-1-0.1 |
| Craves Sugar | 37.83 | 13.2 | 200-50-0.1-200-1-0.1 |
| Dental Decay | 41.5 | 17.64 | 200-50-0.1-200-1-0.1 |
| Depression | 38.07 | 2.42 | 200-50-0.1-200-1-0.1 |
| Diagnosed VitaminD Deficiency | 42.31 | 14.25 | 200-50-0.1-200-1-0.1 |

| Phenotype/ PRSice | AUC | SD | p1-r2-kb-window-size-shiftsize-LDThreshold |
| --- | --- | --- | --- |
| Diagnosed with Sleep Apnea | 47.88 | 26.8 | 200-50-0.1-200-1-0.1 |
| Dimple in the Chin | 41.44 | 9.97 | 200-50-0.1-200-1-0.1 |
| Do you suffer from Agoraphobia | 43.0 | 7.95 | 200-50-0.1-200-1-0.1 |
| Do you grind your teeth | 47.0 | 3.79 | 200-50-0.1-200-1-0.1 |
| Do you have dust mite allergies | 45.96 | 12.36 | 200-50-0.1-200-1-0.1 |
| Dyscalculia | 54.99 | 17.4 | 200-50-0.1-200-1-0.1 |
| Dyslexia | 62.62 | 18.71 | 200-50-0.1-200-1-0.1 |
| Ear Infections In Childhood | 50.54 | 14.63 | 200-50-0.1-200-1-0.1 |
| Ear darwins tubercle | 30.97 | 12.76 | 200-50-0.1-200-1-0.1 |
| Earlobe Free or Attached | 58.62 | 6.53 | 200-50-0.1-200-1-0.1 |
| Ear wiggling | 58.08 | 14.83 | 200-50-0.1-200-1-0.1 |
| Energy Level | 40.89 | 22.49 | 200-50-0.1-200-1-0.1 |
| Enjoy riding a motorbike | 53.56 | 24.34 | 200-50-0.1-200-1-0.1 |
| Fibromyalgia | 14.17 | 17.08 | 200-50-0.1-200-1-0.1 |
| Fish Preference | 36.56 | 5.92 | 200-50-0.1-200-1-0.1 |
| Hair Type | 31.53 | 25.51 | 200-50-0.1-200-1-0.1 |
| Have MECFS | 61.62 | 24.02 | 200-50-0.1-200-1-0.1 |
| Hypermobility | 25.98 | 14.66 | 200-50-0.1-200-1-0.1 |
| Hypertension | 40.0 | 19.9 | 200-50-0.1-200-1-0.1 |
| Hypertriglyceridemia | 41.0 | 19.42 | 200-50-0.1-200-1-0.1 |
| Inflammation | 36.67 | 23.09 | 200-50-0.1-200-1-0.1 |
| Irritable Bowel Syndrome | 56.96 | 21.45 | 200-50-0.1-200-1-0.1 |
| Large tonsil crypts and tonsil stones | 43.56 | 25.12 | 200-50-0.1-200-1-0.1 |
| Mental Disease | 35.9 | 6.17 | 200-50-0.1-200-1-0.1 |
| Migraine | 37.83 | 14.27 | 200-50-0.1-200-1-0.1 |
| Migraine frequency | 36.21 | 23.13 | 200-50-0.1-200-1-0.1 |
| Misophonia | 62.83 | 15.54 | 200-50-0.1-200-1-0.1 |
| Mortons Toe | 57.21 | 7.61 | 200-50-0.1-200-1-0.1 |
| Motion sickness | 23.94 | 0.75 | 200-50-0.1-200-1-0.1 |
| Nickel Dermatitis | 51.03 | 17.47 | 200-50-0.1-200-1-0.1 |
| Panic Disorder | 39.8 | 6.45 | 200-50-0.1-200-1-0.1 |
| Persistent Muscle Pain or Fatigue | 43.87 | 14.84 | 200-50-0.1-200-1-0.1 |
| Photoc Sneez Reflex Photoptarmis | 49.77 | 17.45 | 200-50-0.1-200-1-0.1 |
| Plantar Fasciitis | 58.67 | 20.63 | 200-50-0.1-200-1-0.1 |
| Post traumatic Stress Disorder or PTSD | 31.33 | 3.98 | 200-50-0.1-200-1-0.1 |
| Raynauds | 40.93 | 10.99 | 200-50-0.1-200-1-0.1 |
| Scoliosis | 45.83 | 23.2 | 200-50-0.1-200-1-0.1 |
| Seborrhoeic Dermatitis | 29.0 | 11.76 | 200-50-0.1-200-1-0.1 |
| Sensitivity to Mosquito Bites | 58.5 | 17.08 | 200-50-0.1-200-1-0.1 |
| Sleep Disorders | 54.68 | 8.63 | 200-50-0.1-200-1-0.1 |
| Smell of coffee in urine | 54.19 | 13.55 | 200-50-0.1-200-1-0.1 |
| Sport interest | 28.91 | 12.83 | 200-50-0.1-200-1-0.1 |
| Strabismus | 45.46 | 8.71 | 200-50-0.1-200-1-0.1 |
| Sweat eating spicy food | 56.26 | 10.54 | 200-50-0.1-200-1-0.1 |
| Thyroid Issues Cancer | 29.36 | 12.52 | 200-50-0.1-200-1-0.1 |
| TypeIIDiabetes | 51.66 | 9.31 | 200-50-0.1-200-1-0.1 |
| Widows Peak | 63.81 | 19.7 | 200-50-0.1-200-1-0.1 |
| Apthous in mouth tendency | 58.38 | 23.85 | 200-50-0.1-200-1-0.1 |
| Chronically sore neck glands | 51.83 | 19.64 | 200-50-0.1-200-1-0.1 |
| Cluster headache | 46.6 | 6.29 | 200-50-0.1-200-1-0.1 |
| Double jointed thumbs | 81.88 | 11.44 | 200-50-0.1-200-1-0.1 |
| Eczema | 46.67 | 24.72 | 200-50-0.1-200-1-0.1 |
| Generalizedanxietydisorder | 84.67 | 10.63 | 200-50-0.1-200-1-0.1 |
| Hairnear | 49.02 | 9.33 | 200-50-0.1-200-1-0.1 |
| Naturalskinny | 41.81 | 6.18 | 200-50-0.1-200-1-0.1 |
| Restless leg syndrome | 52.0 | 9.89 | 200-50-0.1-200-1-0.1 |
| Ring finger longer than index finger | 61.66 | 0.33 | 200-50-0.1-200-1-0.1 |
| Rolled tongue | 47.61 | 22.16 | 200-50-0.1-200-1-0.1 |
| Wide feet yes no | 51.67 | 40.14 | 200-50-0.1-200-1-0.1 |

| Phenotype/ PRSice | AUC | SD | p1-r2-kb-window-size-shiftsize-LDThreshold |
| --- | --- | --- | --- |

**Table 6:**
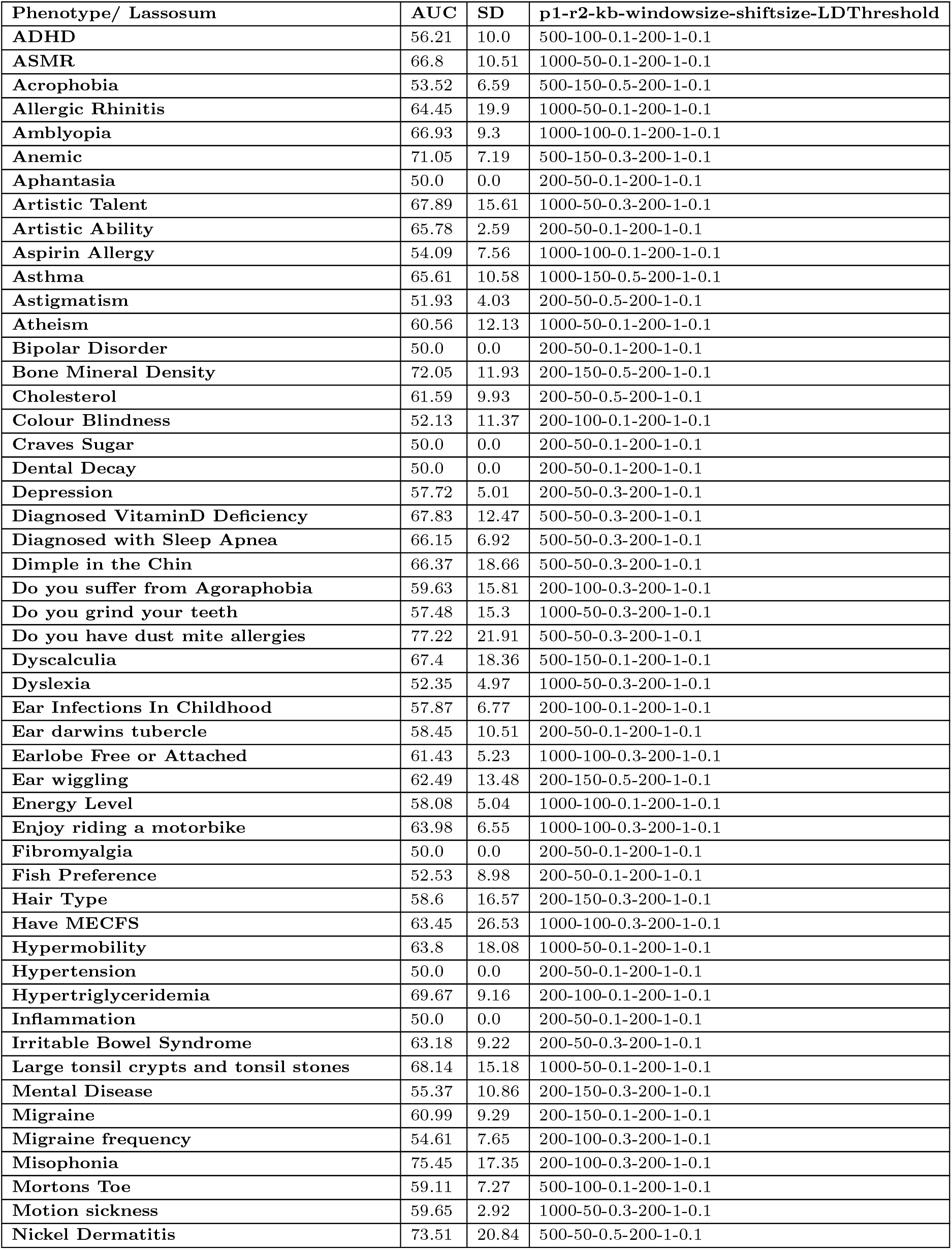
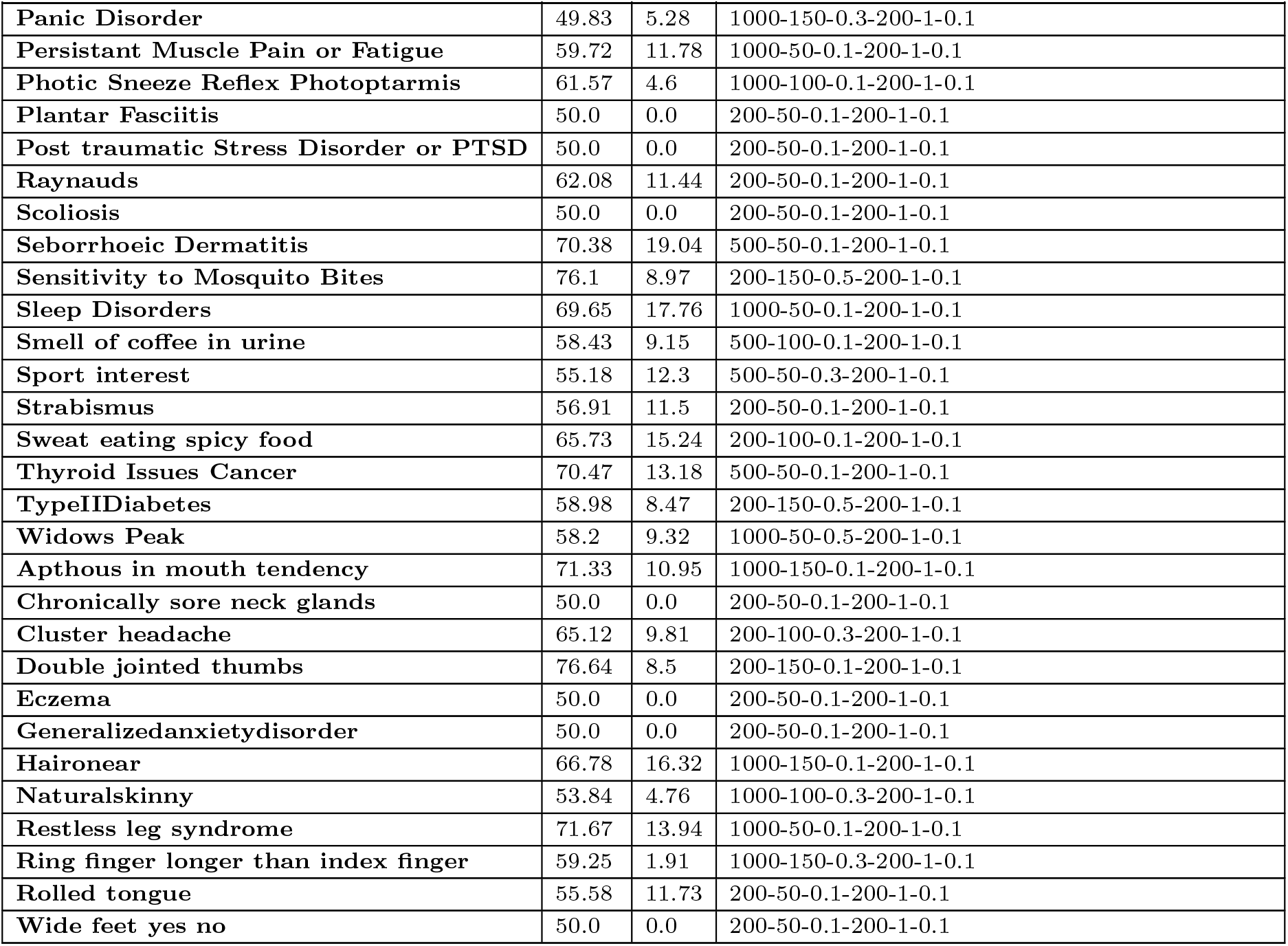
This table shows the Lassosum results. The first column reports the phenotype, the second column shows the best AUC, the third column shows the standard deviation and the fourth column shows the hyperparameters or predictors that generated the best AUC among all other models and hyperparameters.

**Table 7:**
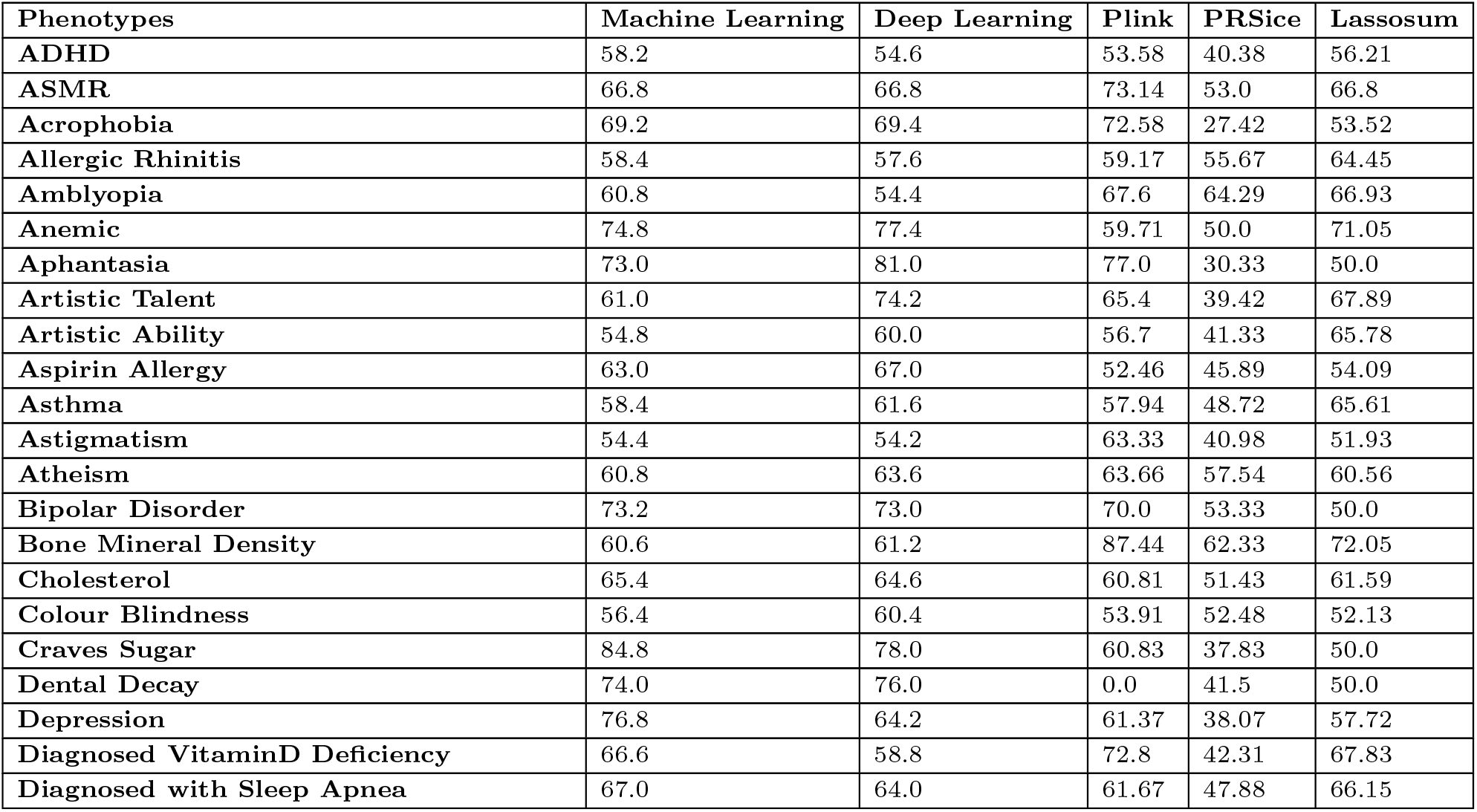
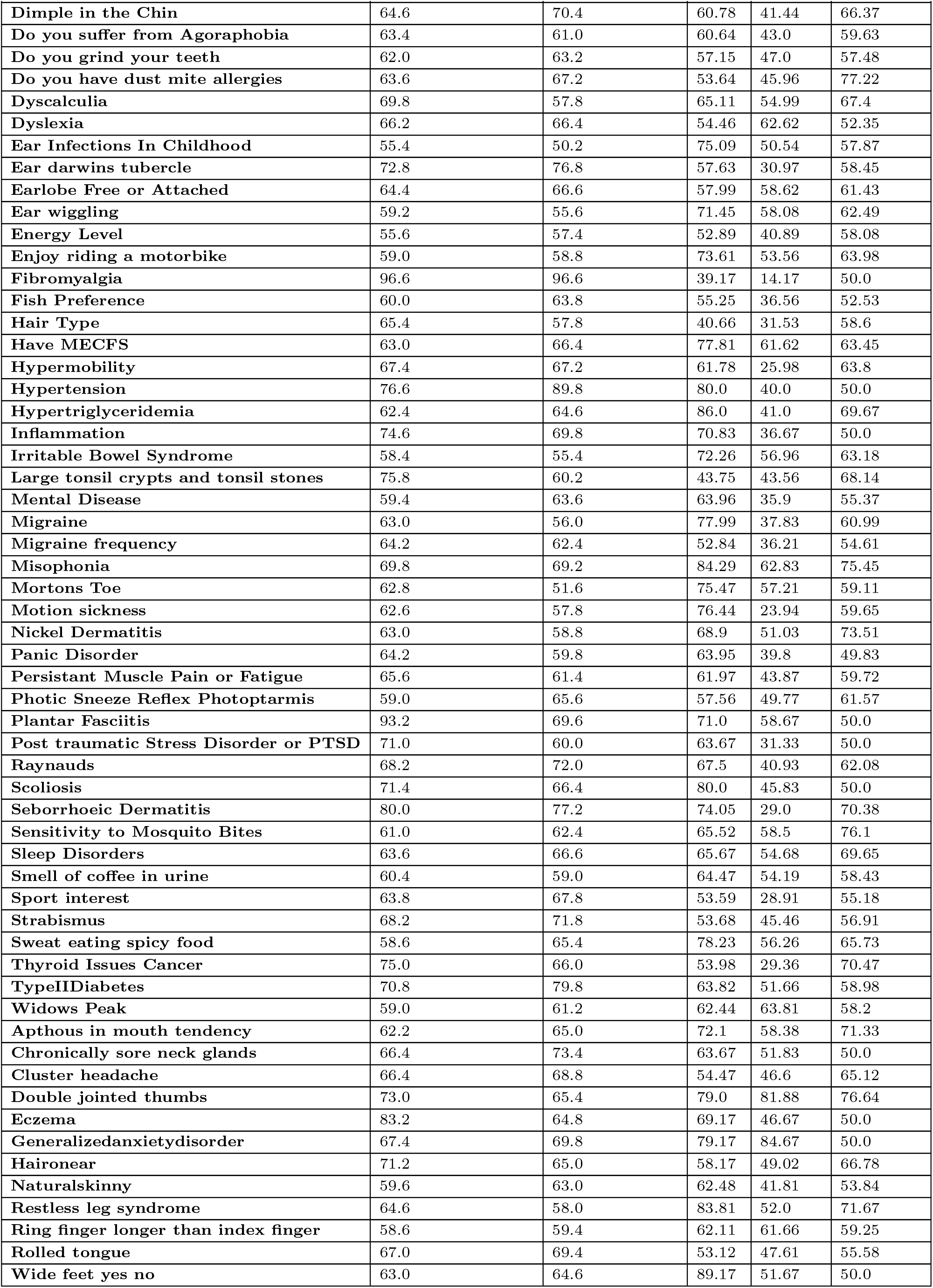
A table reporting the best AUC achieved by each tool for all phenotypes.

| Phenotypes | Machine Learning | Deep Learning | Plink | PRSice | Lassosum |
| --- | --- | --- | --- | --- | --- |
| ADHD | 58.2 | 54.6 | 53.58 | 40.38 | 56.21 |
| ASMR | 66.8 | 66.8 | 73.14 | 53.0 | 66.8 |
| Acrophobia | 69.2 | 69.4 | 72.58 | 27.42 | 53.52 |
| Allergic Rhinitis | 58.4 | 57.6 | 59.17 | 55.67 | 64.45 |
| Amblyopia | 60.8 | 54.4 | 67.6 | 64.29 | 66.93 |
| Anemic | 74.8 | 77.4 | 59.71 | 50.0 | 71.05 |
| Aphantasia | 73.0 | 81.0 | 77.0 | 30.33 | 50.0 |
| Artistic Talent | 61.0 | 74.2 | 65.4 | 39.42 | 67.89 |
| Artistic Ability | 54.8 | 60.0 | 56.7 | 41.33 | 65.78 |
| Aspirin Allergy | 63.0 | 67.0 | 52.46 | 45.89 | 54.09 |
| Asthma | 58.4 | 61.6 | 57.94 | 48.72 | 65.61 |
| Astigmatism | 54.4 | 54.2 | 63.33 | 40.98 | 51.93 |
| Atheism | 60.8 | 63.6 | 63.66 | 57.54 | 60.56 |
| Bipolar Disorder | 73.2 | 73.0 | 70.0 | 53.33 | 50.0 |
| Bone Mineral Density | 60.6 | 61.2 | 87.44 | 62.33 | 72.05 |
| Cholesterol | 65.4 | 64.6 | 60.81 | 51.43 | 61.59 |
| Colour Blindness | 56.4 | 60.4 | 53.91 | 52.48 | 52.13 |
| Craves Sugar | 84.8 | 78.0 | 60.83 | 37.83 | 50.0 |
| Dental Decay | 74.0 | 76.0 | 0.0 | 41.5 | 50.0 |
| Depression | 76.8 | 64.2 | 61.37 | 38.07 | 57.72 |
| Diagnosed VitaminD Deficiency | 66.6 | 58.8 | 72.8 | 42.31 | 67.83 |
| Diagnosed with Sleep Apnea | 67.0 | 64.0 | 61.67 | 47.88 | 66.15 |
| Dimple in the Chin | 64.6 | 70.4 | 60.78 | 41.44 | 66.37 |
| Do you suffer from Agoraphobia | 63.4 | 61.0 | 60.64 | 43.0 | 59.63 |
| Do you grind your teeth | 62.0 | 63.2 | 57.15 | 47.0 | 57.48 |
| Do you have dust mite allergies | 63.6 | 67.2 | 53.64 | 45.96 | 77.22 |
| Dyscalculia | 69.8 | 57.8 | 65.11 | 54.99 | 67.4 |
| Dyslexia | 66.2 | 66.4 | 54.46 | 62.62 | 52.35 |
| Ear Infections In Childhood | 55.4 | 50.2 | 75.09 | 50.54 | 57.87 |
| Ear darwins tubercle | 72.8 | 76.8 | 57.63 | 30.97 | 58.45 |
| Earlobe Free or Attached | 64.4 | 66.6 | 57.99 | 58.62 | 61.43 |
| Ear wiggling | 59.2 | 55.6 | 71.45 | 58.08 | 62.49 |
| Energy Level | 55.6 | 57.4 | 52.89 | 40.89 | 58.08 |
| Enjoy riding a motorbike | 59.0 | 58.8 | 73.61 | 53.56 | 63.98 |
| Fibromyalgia | 96.6 | 96.6 | 39.17 | 14.17 | 50.0 |
| Fish Preference | 60.0 | 63.8 | 55.25 | 36.56 | 52.53 |
| Hair Type | 65.4 | 57.8 | 40.66 | 31.53 | 58.6 |
| Have MECFS | 63.0 | 66.4 | 77.81 | 61.62 | 63.45 |
| Hypermobility | 67.4 | 67.2 | 61.78 | 25.98 | 63.8 |
| Hypertension | 76.6 | 89.8 | 80.0 | 40.0 | 50.0 |
| Hypertriglyceridemia | 62.4 | 64.6 | 86.0 | 41.0 | 69.67 |
| Inflammation | 74.6 | 69.8 | 70.83 | 36.67 | 50.0 |
| Irritable Bowel Syndrome | 58.4 | 55.4 | 72.26 | 56.96 | 63.18 |
| Large tonsil crypts and tonsil stones | 75.8 | 60.2 | 43.75 | 43.56 | 68.14 |
| Mental Disease | 59.4 | 63.6 | 63.96 | 35.9 | 55.37 |
| Migraine | 63.0 | 56.0 | 77.99 | 37.83 | 60.99 |
| Migraine frequency | 64.2 | 62.4 | 52.84 | 36.21 | 54.61 |
| Misophonia | 69.8 | 69.2 | 84.29 | 62.83 | 75.45 |
| Mortons Toe | 62.8 | 51.6 | 75.47 | 57.21 | 59.11 |
| Motion sickness | 62.6 | 57.8 | 76.44 | 23.94 | 59.65 |
| Nickel Dermatitis | 63.0 | 58.8 | 68.9 | 51.03 | 73.51 |
| Panic Disorder | 64.2 | 59.8 | 63.95 | 39.8 | 49.83 |
| Persistent Muscle Pain or Fatigue | 65.6 | 61.4 | 61.97 | 43.87 | 59.72 |
| Photoc Sneez Reflex Photoptarmis | 59.0 | 65.6 | 57.56 | 49.77 | 61.57 |
| Plantar Fasciitis | 93.2 | 69.6 | 71.0 | 58.67 | 50.0 |
| Post traumatic Stress Disorder or PTSD | 71.0 | 60.0 | 63.67 | 31.33 | 50.0 |
| Raynauds | 68.2 | 72.0 | 67.5 | 40.93 | 62.08 |
| Scoliosis | 71.4 | 66.4 | 80.0 | 45.83 | 50.0 |
| Seborrhoeic Dermatitis | 80.0 | 77.2 | 74.05 | 29.0 | 70.38 |
| Sensitivity to Mosquito Bites | 61.0 | 62.4 | 65.52 | 58.5 | 76.1 |
| Sleep Disorders | 63.6 | 66.6 | 65.67 | 54.68 | 69.65 |
| Smell of coffee in urine | 60.4 | 59.0 | 64.47 | 54.19 | 58.43 |
| Sport interest | 63.8 | 67.8 | 53.59 | 28.91 | 55.18 |
| Strabismus | 68.2 | 71.8 | 53.68 | 45.46 | 56.91 |
| Sweat eating spicy food | 58.6 | 65.4 | 78.23 | 56.26 | 65.73 |
| Thyroid Issues Cancer | 75.0 | 66.0 | 53.98 | 29.36 | 70.47 |
| TypeIIDiabetes | 70.8 | 79.8 | 63.82 | 51.66 | 58.98 |
| Widows Peak | 59.0 | 61.2 | 62.44 | 63.81 | 58.2 |
| Apthous in mouth tendency | 62.2 | 65.0 | 72.1 | 58.38 | 71.33 |
| Chronically sore neck glands | 66.4 | 73.4 | 63.67 | 51.83 | 50.0 |
| Cluster headache | 66.4 | 68.8 | 54.47 | 46.6 | 65.12 |
| Double jointed thumbs | 73.0 | 65.4 | 79.0 | 81.88 | 76.64 |
| Eczema | 83.2 | 64.8 | 69.17 | 46.67 | 50.0 |
| Generalizedanxietydisorder | 67.4 | 69.8 | 79.17 | 84.67 | 50.0 |
| Haironear | 71.2 | 65.0 | 58.17 | 49.02 | 66.78 |
| Natural skinny | 59.6 | 63.0 | 62.48 | 41.81 | 53.84 |
| Restless leg syndrome | 64.6 | 58.0 | 83.81 | 52.0 | 71.67 |
| Ring finger longer than index finger | 58.6 | 59.4 | 62.11 | 61.66 | 59.25 |
| Rolled tongue | 67.0 | 69.4 | 53.12 | 47.61 | 55.58 |
| Wide feet yes no | 63.0 | 64.6 | 89.17 | 51.67 | 50.0 |

| Phenotypes | Machine Learning | Deep Learning | Plink | PRSice | Lassosum |
| --- | --- | --- | --- | --- | --- |

**Table 8:**
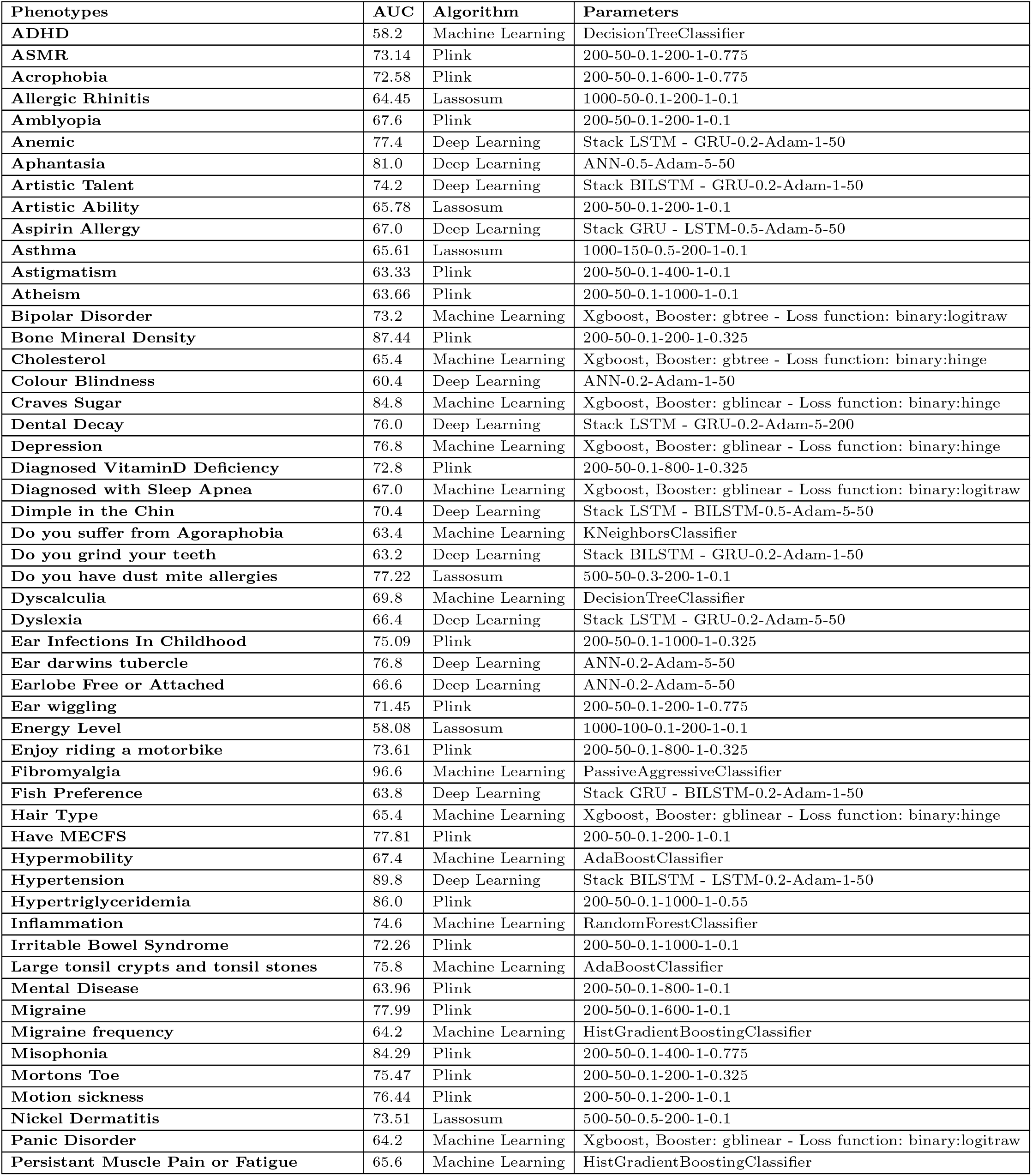
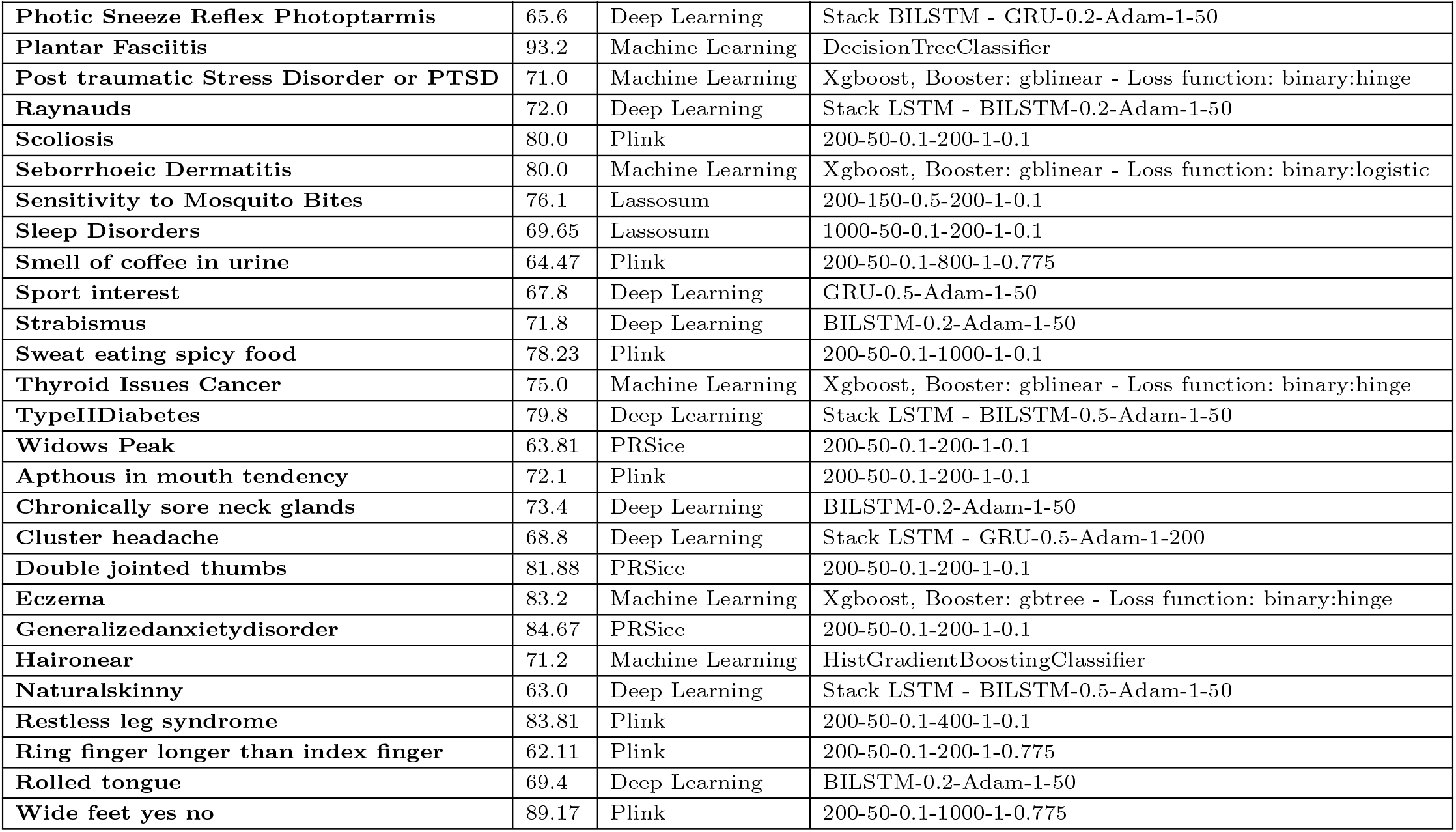
A table showing the best model and corresponding parameters for each phenotype. The first column shows the phenotype, the second column shows the best AUC, the third column shows the algorithm that generated the best AUC, and the third column shows the parameters for the algorithm.

| Phenotypes | AUC | Algorithm | Parameters |
| --- | --- | --- | --- |
| ADHD | 58.2 | Machine Learning | DecisionTreeClassifier |
| ASMR | 73.14 | Plink | 200-50-0.1-200-1-0.775 |
| Acrophobia | 72.58 | Plink | 200-50-0.1-600-1-0.775 |
| Allergic Rhinitis | 64.45 | Lassosum | 1000-50-0.1-200-1-0.1 |
| Amblyopia | 67.6 | Plink | 200-50-0.1-200-1-0.1 |
| Anemic | 77.4 | Deep Learning | Stack LSTM - GRU-0.2-Adam-1-50 |
| Aphantasia | 81.0 | Deep Learning | ANN-0.5-Adam-5-50 |
| Artistic Talent | 74.2 | Deep Learning | Stack BiLSTM - GRU-0.2-Adam-1-50 |
| Artistic Ability | 65.78 | Lassosum | 200-50-0.1-200-1-0.1 |
| Aspirin Allergy | 67.0 | Deep Learning | Stack GRU - LSTM-0.5-Adam-5-50 |
| Asthma | 65.61 | Lassosum | 1000-150-0.5-200-1-0.1 |
| Astigmatism | 63.33 | Plink | 200-50-0.1-400-1-0.1 |
| Atheism | 63.66 | Plink | 200-50-0.1-1000-1-0.1 |
| Bipolar Disorder | 73.2 | Machine Learning | Xgboost, Booster: gbtrees - Loss function: binary:logitraw |
| Bone Mineral Density | 87.44 | Plink | 200-50-0.1-200-1-0.325 |
| Cholesterol | 65.4 | Machine Learning | Xgboost, Booster: gbtrees - Loss function: binary:hinge |
| Colour Blindness | 60.4 | Deep Learning | ANN-0.2-Adam-1-50 |
| Craves Sugar | 84.8 | Machine Learning | Xgboost, Booster: gblinear - Loss function: binary:hinge |
| Dental Decay | 76.0 | Deep Learning | Stack LSTM - GRU-0.2-Adam-5-200 |
| Depression | 76.8 | Machine Learning | Xgboost, Booster: gblinear - Loss function: binary:hinge |
| Diagnosed VitaminD Deficiency | 72.8 | Plink | 200-50-0.1-800-1-0.325 |
| Diagnosed with Sleep Apnea | 67.0 | Machine Learning | Xgboost, Booster: gblinear - Loss function: binary:logitraw |
| Dimple in the Chin | 70.4 | Deep Learning | Stack LSTM - BiLSTM-0.5-Adam-5-50 |
| Do you suffer from Agoraphobia | 63.4 | Machine Learning | KNeighborsClassifier |
| Do you grind your teeth | 63.2 | Deep Learning | Stack BiLSTM - GRU-0.2-Adam-1-50 |
| Do you have dust mite allergies | 77.22 | Lassosum | 500-50-0.3-200-1-0.1 |
| Dyscalculia | 69.8 | Machine Learning | DecisionTreeClassifier |
| Dyslexia | 66.4 | Deep Learning | Stack LSTM - GRU-0.2-Adam-5-50 |
| Ear Infections In Childhood | 75.09 | Plink | 200-50-0.1-1000-1-0.325 |
| Ear darwins tubercle | 76.8 | Deep Learning | ANN-0.2-Adam-5-50 |
| Earlobe Free or Attached | 66.6 | Deep Learning | ANN-0.2-Adam-5-50 |
| Ear wiggling | 71.45 | Plink | 200-50-0.1-200-1-0.775 |
| Energy Level | 58.08 | Lassosum | 1000-100-0.1-200-1-0.1 |
| Enjoy riding a motorbike | 73.61 | Plink | 200-50-0.1-800-1-0.325 |
| Fibromyalgia | 96.6 | Machine Learning | PassiveAggressiveClassifier |
| Fish Preference | 63.8 | Deep Learning | Stack GRU - BiLSTM-0.2-Adam-1-50 |
| Hair Type | 65.4 | Machine Learning | Xgboost, Booster: gblinear - Loss function: binary:hinge |
| Have MECFS | 77.81 | Plink | 200-50-0.1-200-1-0.1 |
| Hypermobility | 67.4 | Machine Learning | AdaBoostClassifier |
| Hypertension | 89.8 | Deep Learning | Stack BiLSTM - LSTM-0.2-Adam-1-50 |
| Hypertriglyceridemia | 86.0 | Plink | 200-50-0.1-1000-1-0.55 |
| Inflammation | 74.6 | Machine Learning | RandomForestClassifier |
| Irritable Bowel Syndrome | 72.26 | Plink | 200-50-0.1-1000-1-0.1 |
| Large tonsil crypts and tonsil stones | 75.8 | Machine Learning | AdaBoostClassifier |
| Mental Disease | 63.96 | Plink | 200-50-0.1-800-1-0.1 |
| Migraine | 77.99 | Plink | 200-50-0.1-600-1-0.1 |
| Migraine frequency | 64.2 | Machine Learning | HistGradientBoostingClassifier |
| Misophonia | 84.29 | Plink | 200-50-0.1-400-1-0.775 |
| Mortons Toe | 75.47 | Plink | 200-50-0.1-200-1-0.325 |
| Motion sickness | 76.44 | Plink | 200-50-0.1-200-1-0.1 |
| Nickel Dermatitis | 73.51 | Lassosum | 500-50-0.5-200-1-0.1 |
| Panic Disorder | 64.2 | Machine Learning | Xgboost, Booster: gblinear - Loss function: binary:logitraw |
| Persistent Muscle Pain or Fatigue | 65.6 | Machine Learning | HistGradientBoostingClassifier |
| Photoc Sneeze Reflex Photoptarmis | 65.6 | Deep Learning | Stack BILSTM - GRU-0.2-Adam-1-50 |
| Plantar Fasciitis | 93.2 | Machine Learning | DecisionTreeClassifier |
| Post traumatic Stress Disorder or PTSD | 71.0 | Machine Learning | Xgboost, Booster: gblinear - Loss function: binary:hinge |
| Raynauds | 72.0 | Deep Learning | Stack LSTM - BILSTM-0.2-Adam-1-50 |
| Scoliosis | 80.0 | Plink | 200-50-0.1-200-1-0.1 |
| Seborrhoeic Dermatitis | 80.0 | Machine Learning | Xgboost, Booster: gblinear - Loss function: binary:logistic |
| Sensitivity to Mosquito Bites | 76.1 | Lassosum | 200-150-0.5-200-1-0.1 |
| Sleep Disorders | 69.65 | Lassosum | 1000-50-0.1-200-1-0.1 |
| Smell of coffee in urine | 64.47 | Plink | 200-50-0.1-800-1-0.775 |
| Sport interest | 67.8 | Deep Learning | GRU-0.5-Adam-1-50 |
| Strabismus | 71.8 | Deep Learning | BILSTM-0.2-Adam-1-50 |
| Sweat eating spicy food | 78.23 | Plink | 200-50-0.1-1000-1-0.1 |
| Thyroid Issues Cancer | 75.0 | Machine Learning | Xgboost, Booster: gblinear - Loss function: binary:hinge |
| TypeIIDiabetes | 79.8 | Deep Learning | Stack LSTM - BILSTM-0.5-Adam-1-50 |
| Widows Peak | 63.81 | PRSize | 200-50-0.1-200-1-0.1 |
| Apthous in mouth tendency | 72.1 | Plink | 200-50-0.1-200-1-0.1 |
| Chronically sore neck glands | 73.4 | Deep Learning | BILSTM-0.2-Adam-1-50 |
| Cluster headache | 68.8 | Deep Learning | Stack LSTM - GRU-0.5-Adam-1-200 |
| Double jointed thumbs | 81.88 | PRSize | 200-50-0.1-200-1-0.1 |
| Eczema | 83.2 | Machine Learning | Xgboost, Booster: gbtrees - Loss function: binary:hinge |
| Generalizedanxietydisorder | 84.67 | PRSize | 200-50-0.1-200-1-0.1 |
| Haironear | 71.2 | Machine Learning | HistGradientBoostingClassifier |
| Naturalskinny | 63.0 | Deep Learning | Stack LSTM - BILSTM-0.5-Adam-1-50 |
| Restless leg syndrome | 83.81 | Plink | 200-50-0.1-400-1-0.1 |
| Ring finger longer than index finger | 62.11 | Plink | 200-50-0.1-200-1-0.775 |
| Rolled tongue | 69.4 | Deep Learning | BILSTM-0.2-Adam-1-50 |
| Wide feet yes no | 89.17 | Plink | 200-50-0.1-1000-1-0.775 |

## References

1. Samuel J. Virolainen, Andrew VonHandorf, Kenyatta C. M. F. Viel, Matthew T. Weirauch, and Leah C. Kottyan. Gene–environment interactions and their impact on human health. Genes and Immunity, 24(1):1–11, December 2022.

2. Danielle M. Dick. Gene-environment interaction in psychological traits and disorders. Annual Review of Clinical Psychology, 7(1):383–409, April 2011.

3. Roxana Daneshjou, Yanran Wang, Yana Bromberg, Samuele Bovo, Pier L Martelli, Giulia Babbi, Pietro Di Lena, Rita Casadio, Matthew Edwards, David Gifford, David T Jones, Laksshman Sundaram, Rajendra Rana Bhat, Xiaolin Li, Lipika R. Pal, Kunal Kundu, Yizhou Yin, John Moult, Yuxiang Jiang, Vikas Pejaver, Kymberleigh A. Pagel, Biao Li, Sean D. Mooney, Predrag Radivojac, Sohela Shah, Marco Carraro, Alessandra Gasparini, Emanuela Leonardi, Manuel Giollo, Carlo Ferrari, Silvio C E Tosatto, Eran Bachar, Johnathan R. Azaria, Yanay Ofran, Ron Unger, Abhishek Niroula, Mauno Vihinen, Billy Chang, Maggie H Wang, Andre Franke, Britt-Sabina Petersen, Mehdi Pirooznia, Peter Zandi, Richard McCombie, James B. Potash, Russ B. Altman, Teri E. Klein, Roger A. Hoskins, Susanna Repo, Steven E. Brenner, and Alexander A. Morgan. Working toward precision medicine: Predicting phenotypes from exomes in the critical assessment of genome interpretation (CAGI) challenges. Human Mutation, 38(9):1182–1192, July 2017.

4. Xinran Dong, Tiantian Xiao, Bin Chen, Yulan Lu, and Wenhao Zhou. Precision medicine via the integration of phenotype-genotype information in neonatal genome project. Fundamental Research, 2(6):873–884, November 2022.

5. Chee Seng Ku, En Yun Loy, Agus Salim, Yudi Pawitan, and Kee Seng Chia. The discovery of human genetic variations and their use as disease markers: past, present and future. Journal of Human Genetics, 55(7):403–415, May 2010.

6. S O Y Keita, R A Kittles, C D M Royal, G E Bonney, P Furbert-Harris, G M Dunston, and C N Rotimi. Conceptualizing human variation. Nature Genetics, 36(S11):S17–S20, October 2004.

7. Emil Uffelmann, Qin Qin Huang, Nchangwi Syntia Munung, Jantina de Vries, Yukinori Okada, Alicia R. Martin, Hilary C. Martin, Tuuli Lappalainen, and Danielle Posthuma. Genome-wide association studies. Nature Reviews Methods Primers, 1(1), August 2021.

8. J.R. Shaffer, E. Feingold, and M.L. Marazita. Genomewide association studies. Journal of Dental Research, 91(7):637–641, May 2012.

9. Vivian Tam, Nikunj Patel, Michelle Turcotte, Yohan Bossé, Guillaume Paré, and David Meyre. Benefits and limitations of genome-wide association studies. Nature Reviews Genetics, 20(8):467–484, May 2019.

10. Shing Wan Choi, Timothy Shin-Heng Mak, and Paul F. O’Reilly. Tutorial: a guide to performing polygenic risk score analyses. Nature Protocols, 15(9):2759–2772, July 2020.

11. Cathryn M. Lewis and Evangelos Vassos. Polygenic risk scores: from research tools to clinical instruments. Genome Medicine, 12(1), May 2020.

12. Jennifer A. Collister, Xiaonan Liu, and Lei Clifton. Calculating polygenic risk scores (PRS) in UK biobank: A practical guide for epidemiologists. Frontiers in Genetics, 13, February 2022.

13. Jordi Merino, Marta Guasch-Ferré, Jun Li, Wonil Chung, Yang Hu, Baoshan Ma, Yanping Li, Jae H. Kang, Peter Kraft, Liming Liang, Qi Sun, Paul W. Franks, JoAnn E. Manson, Walter C. Willet, Jose C. Florez, and Frank B. Hu. Polygenic scores, diet quality, and type 2 diabetes risk: An observational study among 35, 759 adults from 3 US cohorts. PLOS Medicine, 19(4):e1003972, April 2022.

14. Ebba Du Rietz, Jonathan Coleman, Kylie Glanville, Shing Wan Choi, Paul F. O’Reilly, and Jonna Kuntsi. Association of polygenic risk for attention-deficit/hyperactivity disorder with co-occurring traits and disorders. Biological Psychiatry: Cognitive Neuroscience and Neuroimaging, 3(7):635–643, July 2018.

15. Jack W. O’Sullivan, Sridharan Raghavan, Carla Marquez-Luna, Jasmine A. Luzum, Scott M. Damrauer, Euan A. Ashley, Christopher J. O’Donnell, Cristen J. Willer, and Pradeep Natarajan and. Polygenic risk scores for cardiovascular disease: A scientific statement from the american heart association. Circulation, 146(8), August 2022.

16. Daniele Raimondi, Gabriele Orlando, Nora Verplaetse, Piero Fariselli, and Yves Moreau. Editorial: Towards genome interpretation: Computational methods to model the genotype-phenotype relationship. Frontiers in Bioinformatics, 2, November 2022.

17. Peter D. Tonner, Abe Pressman, and David Ross. Interpretable modeling of genotype–phenotype landscapes with state-of-the-art predictive power. Proceedings of the National Academy of Sciences, 119(26), June 2022.

18. Douglas E. V. Pires, Jing Chen, Tom L. Blundell, and David B. Ascher. In silico functional dissection of saturation mutagenesis: Interpreting the relationship between phenotypes and changes in protein stability, interactions and activity. Scientific Reports, 6(1), January 2016.

19. Monica F. Danilevicz, Mitchell Gill, Robyn Anderson, Jacqueline Batley, Mohammed Bennamoun, Philipp E. Bayer, and David Edwards. Plant genotype to phenotype prediction using machine learning. Frontiers in Genetics, 13, May 2022.

20. Muhammad Muneeb, Samuel Feng, and Andreas Henschel. An empirical comparison between polygenic risk scores and machine learning for case/control classification. February 2022.

21. Tingting Guo and Xianran Li. Machine learning for predicting phenotype from genotype and environment. Current Opinion in Biotechnology, 79:102853, February 2023.

22. Muhammad Muneeb, Samuel F. Feng, and Andreas Henschel. Can we convert genotype sequences into images for cases/controls classification? Frontiers in Bioinformatics, 2, June 2022.

23. João Fadista, Alisa K Manning, Jose C Florez, and Leif Groop. The (in)famous GWAS p-value threshold revisited and updated for low-frequency variants. European Journal of Human Genetics, 24(8):1202–1205, January 2016.

24. Avjinder S. Kaler and Larry C. Purcell. Estimation of a significance threshold for genome-wide association studies. BMC Genomics, 20(1), July 2019.

25. Daniel R Kick, Jason G Wallace, James C Schnable, Judith M Kolkman, Barış Alaca, Timothy M Beissinger, Jode Edwards, David Ertl, Sherry Flint-Garcia, Joseph L Gage, Candice N Hirsch, Joseph E Knoll, Natalia de Leon, Dayane C Lima, Danilo E Moreta, Maninder P Singh, Addie Thompson, Teclemariam Weldekidan, and Jacob D Washburn. Yield prediction through integration of genetic, environment, and management data through deep learning. G3 Genes|Genomes|Genetics, 13(4), January 2023.

26. Nikoletta Katsaouni, Araek Tashkandi, Lena Wiese, and Marcel H. Schulz. Machine learning based disease prediction from genotype data. Biological Chemistry, 402(8):871–885, July 2021.

27. David O. Enoma, Janet Bishung, Theresa Abiodun, Olubanke Ogunlana, and Victor Chukwudi Osamor. Machine learning approaches to genome-wide association studies. Journal of King Saud University - Science, 34(4):101847, June 2022.

28. Sebastian Okser, Tapio Pahikkala, Antti Airola, Tapio Salakoski, Samuli Ripatti, and Tero Aittokallio. Regularized machine learning in the genetic prediction of complex traits. PLoS genetics, 10(11):e1004754, 2014.

29. Sarah Taghavi Namin, Mohammad Esmaeilzadeh, Mohammad Najafi, Tim B Brown, and Justin O Borevitz. Deep phenotyping: deep learning for temporal phenotype/genotype classification. Plant methods, 14(1):1–14, 2018.

30. Yaron Gurovich, Yair Hanani, Omri Bar, Guy Nadav, Nicole Fleischer, Dekel Gelbman, Lina Basel-Salmon, Peter M. Krawitz, Susanne B. Kamphausen, Martin Zenker, Lynne M. Bird, and Karen W. Gripp. Identifying facial phenotypes of genetic disorders using deep learning. Nature Medicine, 25(1):60–64, January 2019.

31. Hye-Young Yoo, Ki-Chan Lee, Ji-Eun Woo, Sung-Ha Park, Sunghoon Lee, Joungsu Joo, Jin-Sik Bae, Hyuk-Jung Kwon, and Byoung-Jun Park. A genome-wide association study and machine-learning algorithm analysis on the prediction of facial phenotypes by genotypes in korean women. Clinical, Cosmetic and Investigational Dermatology, 15:433–445, 2022. PMID: 35313536.

32. Matthew Bracher-Smith, Karen Crawford, and Valentina Escott-Price. Machine learning for genetic prediction of psychiatric disorders: a systematic review. Molecular Psychiatry, 26(1):70–79, June 2020.

33. Jon McCormack and Andy Lomas. Deep learning of individual aesthetics. Neural Computing and Applications, 33(1):3–17, October 2020.

34. Nastasiya F. Grinberg, Oghenejokpeme I. Orhobor, and Ross D. King. An evaluation of machine-learning for predicting phenotype: studies in yeast, rice, and wheat. Machine Learning, 109(2):251–277, October 2019.

35. Wenlong Ma, Zhixu Qiu, Jie Song, Jiajia Li, Qian Cheng, Jingjing Zhai, and Chuang Ma. A deep convolutional neural network approach for predicting phenotypes from genotypes. Planta, 248(5):1307–1318, August 2018.

36. Johnathon Shook, Tryambak Gangopadhyay, Linjiang Wu, Baskar Ganapathysubramanian, Soumik Sarkar, and Asheesh K. Singh. Crop yield prediction integrating genotype and weather variables using deep learning. PLOS ONE, 16(6):e0252402, June 2021.

37. Arti Singh, Baskar Ganapathysubramanian, Asheesh Kumar Singh, and Soumik Sarkar. Machine learning for high-throughput stress phenotyping in plants. Trends in Plant Science, 21(2):110–124, February 2016.

38. Mitchell Gill, Robyn Anderson, Haifei Hu, Mohammed Bennamoun, Jakob Petereit, Babu Valliyodan, Henry T. Nguyen, Jacqueline Batley, Philipp E. Bayer, and David Edwards. Machine learning models outperform deep learning models, provide interpretation and facilitate feature selection for soybean trait prediction. BMC Plant Biology, 22(1), April 2022.

39. Ross KK Leung, Ying Wang, Ronald CW Ma, Andrea OY Luk, Vincent Lam, Maggie Ng, Wing Yee So, Stephen KW Tsui, and Juliana CN Chan. Using a multi-staged strategy based on machine learning and mathematical modeling to predict genotype-phenotype risk patterns in diabetic kidney disease: a prospective case–control cohort analysis. BMC Nephrology, 14(1), July 2013.

40. Giorgio Guzzetta, Giuseppe Jurman, and Cesare Furlanello. A machine learning pipeline for quantitative phenotype prediction from genotype data. BMC Bioinformatics, 11(S8), October 2010.

41. Yang Liu, Duolin Wang, Fei He, Juexin Wang, Trupti Joshi, and Dong Xu. Phenotype prediction and genome-wide association study using deep convolutional neural network of soybean. Frontiers in Genetics, 10, November 2019.

42. Rostam Abdollahi-Arpanahi, Daniel Gianola, and Francisco Penãgaricano. Deep learning versus parametric and ensemble methods for genomic prediction of complex phenotypes. Genetics Selection Evolution, 52(1), February 2020.

43. Joverlyn Gaudillo, Jae Joseph Russell Rodriguez, Allen Nazareno, Lei Rigi Baltazar, Julianne Vilela, Rommel Bulalacao, Mario Domingo, and Jason Albia. Machine learning approach to single nucleotide polymorphism-based asthma prediction. PLOS ONE, 14(12):e0225574, December 2019.

44. Neeraj Budhlakoti, Amar Kant Kushwaha, Anil Rai, K K Chaturvedi, Anuj Kumar, Anjan Kumar Pradhan, Uttam Kumar, Rajeev Ranjan Kumar, Philomin Juliana, D C Mishra, and Sundeep Kumar. Genomic selection: A tool for accelerating the efficiency of molecular breeding for development of climate-resilient crops. Frontiers in Genetics, 13, February 2022.

45. Xinlei Wang, Donghua Li, Sufang Song, Yanhua Zhang, Yuanfang Li, Xiangnan Wang, Danli Liu, Chenxi Zhang, Yanfang Cao, Yawei Fu, Ruili Han, Wenting Li, Xiaojun Liu, Guirong Sun, Guoxi Li, Yadong Tian, Zhuanjian Li, and Xiangtao Kang. Combined transcriptomics and proteomics forecast analysis for potential genes regulating the columbian plumage color in chickens. PLOS ONE, 14(11):e0210850, November 2019.

46. Lisa Bastarache, Joshua C. Denny, and Dan M. Roden. Phenome-wide association studies. JAMA, 327(1):75, January 2022.

47. Bastian Greshake, Philipp E. Bayer, Helge Rausch, and Julia Reda. openSNP–a crowdsourced web resource for personal genomics. PLoS ONE, 9(3):e89204, March 2014.

48. Olivier Naret, David AA Baranger, Sharada Prasanna Mohanty, Bastian Greshake Tzovaras, Marcel Salathé, and Jacques Fellay and. Phenotype prediction from genome-wide genotyping data: a crowdsourcing experiment. bioRxiv, August 2020.

49. Subrata Saha, Himanshu Narayan Singh, Ahmed Soliman, and Sanguthevar Rajasekaran. A novel computational methodology for GWAS multi-locus analysis based on graph theory and machine learning. medRxiv, October 2021.

50. G. Rajesh, X. Mercilin Raajini, K. Martin Sagayam, and Hien Dang. A statistical approach for high order epistasis interaction detection for prediction of diabetic macular edema. Informatics in Medicine Unlocked, 20:100362, 2020.

51. Lu Zhang, Qiuping Pan, Yue Wang, Xintao Wu, and Xinghua Shi. Bayesian Network Construction and Genotype-Phenotype Inference Using GWAS Statistics. IEEE/ACM Transactions on Computational Biology and Bioinformatics, 16(2):475–489, 2019.

52. Javier de Velasco Oriol, Antonio Martinez-Torteya, Victor Trevino, Israel Alanis, Edgar E. Vallejo, and Jose Gerardo Tamez-Pena. Benchmarking machine learning models for the analysis of genetic data using FRESA.CAD Binary Classification Benchmarking. bioRxiv, 2019.

53. Muhammad Muneeb and Andreas Henschel. Eye-color and type-2 diabetes phenotype prediction from genotype data using deep learning methods. BMC Bioinformatics, 22(1), April 2021.

54. Muhammad Muneeb, Samuel F. Feng, and Andreas Henschel. Heritability, genetic variation, and the number of risk SNPs effect on deep learning and polygenic risk scores AUC. In 2022 14th International Conference on Bioinformatics and Biomedical Technology. ACM, May 2022.

55. Justin L. Cope, Hannes A. Baukmann, Jörn E. Klinger, Charles N. J. Ravarani, Erwin P. Böttinger, Stefan Konigorski, and Marco F. Schmidt. Interaction-based feature selection algorithm outperforms polygenic risk score in predicting parkinson’s disease status. Frontiers in Genetics, 12, October 2021.

56. Adrien Badré, Li Zhang, Wellington Muchero, Justin C. Reynolds, and Chongle Pan. Deep neural network improves the estimation of polygenic risk scores for breast cancer. Journal of Human Genetics, 66(4):359–369, October 2020.

57. Damian Gola, Jeannette Erdmann, Bertram Müller-Myhsok, Heribert Schunkert, and Inke R. König. Polygenic risk scores outperform machine learning methods in predicting coronary artery disease status. Genetic Epidemiology, 44(2):125–138, January 2020.

58. Steven Amadeus, Tjeng Wawan Cenggoro, Arif Budiarto, and Bens Pardamean. A design of polygenic risk model with deep learning for colorectal cancer in multiethnic indonesians. Procedia Computer Science, 179:632–639, 2021.

59. Xiaopu Zhou, Yu Chen, Fanny C. F. Ip, Yuanbing Jiang, Han Cao, Ge Lv, Huan Zhong, Jiahang Chen, Tao Ye, Yuewen Chen, Yulin Zhang, Shuangshuang Ma, Ronnie M. N. Lo, Estella P. S. Tong, Michael W. Weiner, Paul Aisen, Ronald Petersen, Clifford R. Jack, William Jagust, John Q. Trojanowski, Arthur W. Toga, Laurel Beckett, Robert C. Green, Andrew J. Saykin, John Morris, Leslie M. Shaw, Zaven Khachaturian, Greg Sorensen, Lew Kuller, Marcus Raichle, Steven Paul, Peter Davies, Howard Fillit, Franz Hefti, David Holtzman, Marek M. Mesulam, William Potter, Peter Snyder, Adam Schwartz, Tom Montine, Ronald G. Thomas, Michael Donohue, Sarah Walter, Devon Gessert, Tamie Sather, Gus Jiminez, Danielle Harvey, Matthew Bernstein, Paul Thompson, Norbert Schuff, Bret Borowski, Jeff Gunter, Matt Senjem, Prashanthi Vemuri, David Jones, Kejal Kantarci, Chad Ward, Robert A. Koeppe, Norm Foster, Eric M. Reiman, Kewei Chen, Chet Mathis, Susan Landau, Nigel J. Cairns, Erin Householder, Lisa Taylor-Reinwald, Virginia Lee, Magdalena Korecka, Michal Figurski, Karen Crawford, Scott Neu, Tatiana M. Foroud, Steven G. Potkin, Li Shen, Kelley Faber, Sungeun Kim, Kwangsik Nho, Leon Thal, Neil Buckholtz, Marylyn Albert, Richard Frank, John Hsiao, Jeffrey Kaye, Joseph Quinn, Betty Lind, Raina Carter, Sara Dolen, Lon S. Schneider, Sonia Pawluczyk, Mauricio Beccera, Liberty Teodoro, Bryan M. Spann, James Brewer, Helen Vanderswag, Adam Fleisher, Judith L. Heidebrink, Joanne L. Lord, Sara S. Mason, Colleen S. Albers, David Knopman, Kris Johnson, Rachelle S. Doody, Javier Villanueva-Meyer, Munir Chowdhury, Susan Rountree, Mimi Dang, Yaakov Stern, Lawrence S. Honig, Karen L. Bell, Beau Ances, Maria Carroll, Sue Leon, Mark A. Mintun, Stacy Schneider, Angela Oliver, Daniel Marson, Randall Griffith, David Clark, David Geldmacher, John Brockington, Erik Roberson, Hillel Grossman, Effie Mitsis, Leyla de Toledo-Morrell, Raj C. Shah, Ranjan Duara, Daniel Varon, Maria T. Greig, Peggy Roberts, Chiadi Onyike, Daniel D’Agostino, Stephanie Kielb, James E. Galvin, Brittany Cerbone, Christina A. Michel, Henry Rusinek, Mony J. de Leon, Lidia Glodzik, Susan De Santi, P. Murali Doraiswamy, Jeffrey R. Petrella, Terence Z. Wong, Steven E. Arnold, Jason H. Karlawish, David Wolk, Charles D. Smith, Greg Jicha, Peter Hardy, Partha Sinha, Elizabeth Oates, Gary Conrad, Oscar L. Lopez, MaryAnn Oakley, Donna M. Simpson, Anton P. Porsteinsson, Bonnie S. Goldstein, Kim Martin, Kelly M. Makino, M. Saleem Ismail, Connie Brand, Ruth A. Mulnard, Gaby Thai, Catherine McAdams-Ortiz, Kyle Womack, Dana Mathews, Mary Quiceno, Ramon Diaz-Arrastia, Richard King, Myron Weiner, Kristen Martin-Cook, Michael DeVous, Allan I. Levey, James J. Lah, Janet S. Cellar, Jeffrey M. Burns, Heather S. Anderson, Russell H. Swerdlow, Liana Apostolova, Kathleen Tingus, Ellen Woo, Daniel H. S. Silverman, Po H. Lu, George Bartzokis, Neill R. Graff-Radford, Francine Parfitt, Tracy Kendall, Heather Johnson, Martin R. Farlow, Ann Marie Hake, Brandy R. Matthews, Scott Herring, Cynthia Hunt, Christopher H. van Dyck, Richard E. Carson, Martha G. MacAvoy, Howard Chertkow, Howard Bergman, Chris Hosein, Ging-Yuek Robin Hsiung, Howard Feldman, Benita Mudge, Michele Assaly, Charles Bernick, Donna Munic, Andrew Kertesz, John Rogers, Dick Trost, Diana Kerwin, Kristine Lipowski, Chuang-Kuo Wu, Nancy Johnson, Carl Sadowsky, Walter Martinez, Teresa Villena, Raymond Scott Turner, Kathleen Johnson, Brigid Reynolds, Reisa A. Sperling, Keith A. Johnson, Gad Marshall, Meghan Frey, Barton Lane, Allyson Rosen, Jared Tinklenberg, Marwan N. Sabbagh, Christine M. Belden, Sandra A. Jacobson, Sherye A. Sirrel, Neil Kowall, Ronald Killiany, Andrew E. Budson, Alexander Norbash, Patricia Lynn Johnson, Joanne Allard, Alan Lerner, Paula Ogrocki, Leon Hudson, Evan Fletcher, Owen Carmichael, John Olichney, Charles DeCarli, Smita Kittur, Michael Borrie, T-Y. Lee, Rob Bartha, Sterling Johnson, Sanjay Asthana, Cynthia M. Carlsson, Adrian Preda, Dana Nguyen, Pierre Tariot, Stephanie Reeder, Vernice Bates, Horacio Capote, Michelle Rainka, Douglas W. Scharre, Maria Kataki, Anahita Adeli, Earl A. Zimmerman, Dzintra Celmins, Alice D. Brown, Godfrey D. Pearlson, Karen Blank, Karen Anderson, Robert B. Santulli, Tamar J. Kitzmiller, Eben S. Schwartz, Kaycee M. Sink, Jeff D. Williamson, Pradeep Garg, Franklin Watkins, Brian R. Ott, Henry Querfurth, Geoffrey Tremont, Stephen Salloway, Paul Malloy, Stephen Correia, Howard J. Rosen, Bruce L. Miller, Jacobo Mintzer, Kenneth Spicer, David Bachman, Stephen Pasternak, Irina Rachinsky, Dick Drost, Nunzio Pomara, Raymundo Hernando, Antero Sarrael, Susan K. Schultz, Laura L. Boles Ponto, Hyungsub Shim, Karen Elizabeth Smith, Norman Relkin, Gloria Chaing, Lisa Raudin, Amanda Smith, Kristin Fargher, Balebail Ashok Raj, Thomas Neylan, Jordan Grafman, Melissa Davis, Rosemary Morrison, Jacqueline Hayes, Shannon Finley, Karl Friedl, Debra Fleischman, Konstantinos Arfanakis, Olga James, Dino Massoglia, J. Jay Fruehling, Sandra Harding, Elaine R. Peskind, Eric C. Petrie, Gail Li, Jerome A. Yesavage, Joy L. Taylor, Ansgar J. Furst, Vincent C. T. Mok, Timothy C. Y. Kwok, Qihao Guo, Kin Y. Mok, Maryam Shoai, John Hardy, Lei Chen, Amy K. Y. Fu, and Nancy Y. Ip and. Deep learning-based polygenic risk analysis for alzheimer’s disease prediction. Communications Medicine, 3(1), April 2023.

60. Jiajie Peng, Jingyi Li, Ruijiang Han, Yuxian Wang, Lu Han, Jinghao Peng, Tao Wang, Jianye Hao, Xuequn Shang, and Zhongyu Wei. A deep learning-based genome-wide polygenic risk score for common diseases identifies individuals with risk. November 2021.

61. Shivani Sehrawat, Keyhan Najafian, and Lingling Jin. Predicting phenotypes from novel genomic markers using deep learning. Bioinformatics Advances, 3(1), January 2023.

62. Bradley Monk, Andrei Rajkovic, Semar Petrus, Aleks Rajkovic, Terry Gaasterland, and Roberto Malinow. A machine learning method to identify genetic variants potentially associated with alzheimer’s disease. Frontiers in Genetics, 12, June 2021.

63. Ying Ma and Xiang Zhou. Genetic prediction of complex traits with polygenic scores: a statistical review. Trends in Genetics, 37(11):995–1011, November 2021.

64. Sinead English, Ido Pen, Nicholas Shea, and Tobias Uller. The information value of non-genetic inheritance in plants and animals. PLOS ONE, 10(1):e0116996, January 2015.

65. Shuangning Li, Zhimei Ren, Chiara Sabatti, and Matteo Sesia. Transfer learning in genome-wide association studies with knockoffs. Sankhya B, November 2022.

66. Muhammad Muneeb, Samuel Feng, and Andreas Henschel. Transfer learning for genotype–phenotype prediction using deep learning models. BMC Bioinformatics, 23(1), November 2022.

67. Florian Privé, Bjarni J. Vilhjálmsson, Hugues Aschard, and Michael G.B. Blum. Making the most of clumping and thresholding for polygenic scores. The American Journal of Human Genetics, 105(6):1213–1221, December 2019.

68. Santiago Alvarez Prado, Isabelle Sanchez, Llorenç Cabrera-Bosquet, Antonin Grau, Claude Welcker, François Tardieu, and Nadine Hilgert. To clean or not to clean phenotypic datasets for outlier plants in genetic analyses? Journal of Experimental Botany, 70(15):3693– 3698, April 2019.

69. Muhammad Muneeb, Samuel F. Feng, and Andreas Henschel. Tutorial on 8 genotype files conversion. In 2022 10th International Conference on Bioinformatics and Computational Biology (ICBCB), pages 13–17, 2022.

70. Yagoub Adam, Chaimae Samtal, Jean Tristan Brandenburg, Oluwadamilare Falola, and Ezekiel Adebiyi. Performing post-genome-wide association study analysis: overview, challenges and recommendations. F1000Research, 10:1002, October 2021.

71. Wenhan Chen, Yang Wu, Zhili Zheng, Ting Qi, Peter M. Visscher, Zhihong Zhu, and Jian Yang. Improved analyses of GWAS summary statistics by reducing data heterogeneity and errors. Nature Communications, 12(1), December 2021.

72. Ali Al-Fatlawi, Negin Malekian, Sebastián García, Andreas Henschel, Ilwook Kim, Andreas Dahl, Beatrix Jahnke, Peter Bailey, Sarah Naomi Bolz, Anna R Poetsch, et al. Deep learning improves pancreatic cancer diagnosis using rna-based variants. Cancers, 13(11):2654, 2021.

73. Princess P. Silva, Joverlyn D. Gaudillo, Julianne A. Vilela, Ranzivelle Marianne L. Roxas-Villanueva, Beatrice J. Tiangco, Mario R. Domingo, and Jason R. Albia. A machine learning-based SNP-set analysis approach for identifying disease-associated susceptibility loci. Scientific Reports, 12(1), September 2022.

74. Aleksandr Medvedev, Satyarth Mishra Sharma, Evgenii Tsatsorin, Elena Nabieva, and Dmitry Yarotsky. Human genotype-to-phenotype predictions: Boosting accuracy with nonlinear models. PLOS ONE, 17(8):e0273293, August 2022.

75. Lizhi Liao, Heng Li, Weiyi Shang, and Lei Ma. An empirical study of the impact of hyperparameter tuning and model optimization on the performance properties of deep neural networks. ACM Transactions on Software Engineering and Methodology, 31(3):1–40, April 2022.

76. J. Crossa. Methodologies for estimating the sample size required for genetic conservation of outbreeding crops. Theoretical and Applied Genetics, 77(2):153–161, February 1989.

77. Wan-Mei Qu, Ni Liang, Zi-Ku Wu, You-Gang Zhao, and Dong Chu. Minimum sample sizes for invasion genomics: Empirical investigation in an invasive whitefly. Ecology and Evolution, 10(1):38–49, October 2019.

78. Carl A Anderson, Fredrik H Pettersson, Geraldine M Clarke, Lon R Cardon, Andrew P Morris, and Krina T Zondervan. Data quality control in genetic case-control association studies. Nature Protocols, 5(9):1564–1573, August 2010.

79. Cathy C. Laurie, Kimberly F. Doheny, Daniel B. Mirel, Elizabeth W. Pugh, Laura J. Bierut, Tushar Bhangale, Frederick Boehm, Neil E. Caporaso, Marilyn C. Cornelis, Howard J. Edenberg, Stacy B. Gabriel, Emily L. Harris, Frank B. Hu, Kevin B. Jacobs, Peter Kraft, Maria Teresa Landi, Thomas Lumley, Teri A. Manolio, Caitlin McHugh, Ian Painter, Justin Paschall, John P. Rice, Kenneth M. Rice, Xiuwen Zheng, and Bruce S. Weir and. Quality control and quality assurance in genotypic data for genome-wide association studies. Genetic Epidemiology, 34(6):591–602, August 2010.

80. Amel Lamri, Shihong Mao, Dipika Desai, Milan Gupta, Guillaume Paré, and Sonia S. Anand. Fine-tuning of genome-wide polygenic risk scores and prediction of gestational diabetes in south asian women. Scientific Reports, 0(1), June 2020.

81. F. Pedregosa, G. Varoquaux, A. Gramfort, V. Michel, B. Thirion, O. Grisel, M. Blondel, P. Prettenhofer, R. Weiss, V. Dubourg, J. Vanderplas, A. Passos, D. Cournapeau, M. Brucher, M. Perrot, and E. Duchesnay. Scikit-learn: Machine learning in Python. Journal of Machine Learning Research, 12:2825–2830, 2011.

82. Bahzad Jijo and Adnan Mohsin Abdulazeez. Classification based on decision tree algorithm for machine learning. Journal of Applied Science and Technology Trends, 2:20–28, 01 2021.

83. Tianqi Chen and Carlos Guestrin. XGBoost: A scalable tree boosting system. In Proceedings of the 22nd ACM SIGKDD International Conference on Knowledge Discovery and Data Mining, KDD ‘16, pages 785–794, New York, NY, USA, 2016. ACM.

84. Nello Cristianini and Elisa Ricci. Support vector machines. In Encyclopedia of Algorithms, pages 928–932. Springer US, 2008.

85. Warren S McCulloch and Walter Pitts. A logical calculus of the ideas immanent in nervous activity. The bulletin of mathematical biophysics, 5(4):115–133, 1943.

86. Rahul Dey and Fathi M. Salem. Gate-variants of gated recurrent unit (gru) neural networks. In 2017 IEEE 60th International Midwest Symposium on Circuits and Systems (MWSCAS), pages 1597–1600, 2017.

87. Sepp Hochreiter and Jürgen Schmidhuber. Long shortterm memory. Neural Computation, 9(8):1735–1780, November 1997.

88. M. Schuster and K.K. Paliwal. Bidirectional recurrent neural networks. IEEE Transactions on Signal Processing, 45(11):2673–2681, 1997.

89. Farhad Pouladi, Hojjat Salehinejad, and Amir Mohammad Gilani. Recurrent neural networks for sequential phenotype prediction in genomics. In 2015 International Conference on Developments of E-Systems Engineering (DeSE). IEEE, December 2015.

90. Parvathaneni Naga Srinivasu, Jana Shafi, T Balamurali Krishna, Canavoy Narahari Sujatha, S Phani Praveen, and Muhammad Fazal Ijaz. Using recurrent neural networks for predicting type-2 diabetes from genomic and tabular data. Diagnostics, 12(12):3067, December 2022.

91. Muhammad Uzair and Noreen Jamil. Effects of hidden layers on the efficiency of neural networks. In 2020 IEEE 23rd International Multitopic Conference (INMIC). IEEE, November 2020.

92. Pérez-Enciso and Zingaretti. A guide for using deep learning for complex trait genomic prediction. Genes, 10(7):553, July 2019.

93. Shaun Purcell, Benjamin Neale, Kathe Todd-Brown, Lori Thomas, Manuel A.R. Ferreira, David Bender, Julian Maller, Pamela Sklar, Paul I.W. de Bakker, Mark J. Daly, and Pak C. Sham. PLINK: A tool set for whole-genome association and population-based linkage analyses. The American Journal of Human Genetics, 81(3):559–575, September 2007.

94. Jack Euesden, Cathryn M. Lewis, and Paul F. O’Reilly. PRSice: Polygenic risk score software. Bioinformatics, 31(9):1466–1468, December 2014.

95. Timothy Shin Heng Mak, Robert Milan Porsch, Shing Wan Choi, Xueya Zhou, and Pak Chung Sham. Polygenic scores via penalized regression on summary statistics. Genetic Epidemiology, 41(6):469–480, May 2017.

96. Amit V. Khera, Mark Chaffin, Krishna G. Aragam, Mary E. Haas, Carolina Roselli, Seung Hoan Choi, Pradeep Natarajan, Eric S. Lander, Steven A. Lubitz, Patrick T. Ellinor, and Sekar Kathiresan. Genome-wide polygenic scores for common diseases identify individuals with risk equivalent to monogenic mutations. Nature Genetics, 50(9):1219–1224, August 2018.

97. Naomi R. Wray, Jian Yang, Michael E. Goddard, and Peter M. Visscher. The genetic interpretation of area under the ROC curve in genomic profiling. PLoS Genetics, 6(2):e1000864, February 2010.

98. Laura Fahey, Derek W. Morris, and Pilib Ó Broin. Comparing the XGBoost machine learning algorithm to polygenic scoring for the prediction of intelligence based on genotype data. June 2022.

99. Muhammad Muneeb, Samuel Feng, and Andreas Henschel. Transfer learning for genotype–phenotype prediction using deep learning models. BMC Bioinformatics, 23(1), November 2022.

100. Shuai Zeng, Ziting Mao, Yijie Ren, Duolin Wang, Dong Xu, and Trupti Joshi. G2pdeep: a web-based deep-learning framework for quantitative phenotype prediction and discovery of genomic markers. Nucleic Acids Research, 49(W1):W228–W236, May 2021.

101. Farhad Pouladi, Hojjat Salehinejad, and Amir Mohammad Gilani. Recurrent neural networks for sequential phenotype prediction in genomics. In 2015 International Conference on Developments of E-Systems Engineering (DeSE), pages 225–230, 2015.

102. Andries T. Marees, Hilde de Kluiver, Sven Stringer, Florence Vorspan, Emmanuel Curis, Cynthia Marie-Claire, and Eske M. Derks. A tutorial on conducting genome-wide association studies: Quality control and statistical analysis. International Journal of Methods in Psychiatric Research, 27(2), February 2018.

103. Matti Pirinen, Peter Donnelly, and Chris C A Spencer. Including known covariates can reduce power to detect genetic effects in case-control studies. Nature Genetics, 44(8):848–851, July 2012.

104. Martín Abadi, Ashish Agarwal, Paul Barham, Eugene Brevdo, Zhifeng Chen, Craig Citro, Greg S. Corrado, Andy Davis, Jeffrey Dean, Matthieu Devin, Sanjay Ghemawat, Ian Goodfellow, Andrew Harp, Geoffrey Irving, Michael Isard, Yangqing Jia, Rafal Jozefowicz, Lukasz Kaiser, Manjunath Kudlur, Josh Levenberg, Dandelion Mané, Rajat Monga, Sherry Moore, Derek Murray, Chris Olah, Mike Schuster, Jonathon Shlens, Benoit Steiner, Ilya Sutskever, Kunal Talwar, Paul Tucker, Vincent Vanhoucke, Vijay Vasudevan, Fernanda Viégas, Oriol Vinyals, Pete Warden, Martin Wattenberg, Martin Wicke, Yuan Yu, and Xiaoqiang Zheng. TensorFlow: Large-scale machine learning on heterogeneous systems, 2015. Software available from tensorflow.org.

105. Francois Chollet et al. Keras, 2015.

## References

1. Bastian Greshake, Philipp E. Bayer, Helge Rausch, and Julia Reda. openSNP–a crowdsourced web resource for personal genomics. PLoS ONE, 9(3):e89204, March 2014.

2. Lindsey A. Ho and Ethan M. Lange. Using public control genotype data to increase power and decrease cost of case–control genetic association studies. Human Genetics, 128(6):597–608, September 2010.

3. Muhammad Muneeb, Samuel F. Feng, and Andreas Henschel. Tutorial on 8 genotype files conversion. In 2022 10th International Conference on Bioinformatics and Computational Biology (ICBCB), pages 13–17, 2022.

4. Rasmus Nielsen, Joshua S. Paul, Anders Albrechtsen, and Yun S. Song. Genotype and SNP calling from next-generation sequencing data. Nature Reviews Genetics, 12(6):443–451, May 2011.

5. Santiago Alvarez Prado, Isabelle Sanchez, Llorenç Cabrera-Bosquet, Antonin Grau, Claude Welcker, François Tardieu, and Nadine Hilgert. To clean or not to clean phenotypic datasets for outlier plants in genetic analyses? Journal of Experimental Botany, 70(15):3693–3698, April 2019.

